# When the pen is mightier than the sword: semi-automatic 2 and 3D image labelling

**DOI:** 10.1101/2024.01.15.575658

**Authors:** Réka Hollandi, David Bauer, Akos Diosdi, Bálint Schrettner, Timea Toth, Dominik Hirling, Gábor Hollandi, Maria Harmati, József Molnár, Peter Horvath

**Affiliations:** Synthetic and Systems Biology Unit, Biological Research Centre HUN-REN, 6726 Szeged, Hungary; Single-Cell Technologies Ltd, 6726 Szeged, Hungary; mAIskin AB, Malmö, 213 76 Sweden; Institute of AI for Health, Helmholtz Zentrum München, 85764 Neuherberg, Germany; Institute for Molecular Medicine Finland (FIMM), University of Helsinki, 00014 Helsinki, Finland

**Keywords:** annotation, minimal contour, 3d, spheroid

## Abstract

Data is the driving engine of learning-based algorithms, the creation of which fundamentally determines the performance, accuracy, generalizability and quality of any model or method trained on it. When only skilled or trained personnel can create reliable annotations, assisted software solutions are desirable to reduce the time and effort the expert must spend on labelling. Herein is proposed an automated annotation helper software package in napari that offers multiple methods to assist the annotator in creating object-based labels on 2D or 3D images.

## Main text

Supervised learning relies on labelled data as its training set. Several clever augmentation techniques have been applied [1-4] to address the problem of inadequate amount of labelled data, even though semi- or self-supervised algorithms with more moderate requirements are also applied in the domain of microscopy image analysis. The most commonly used deep learning-based methods for cell or nucleus segmentation such as StarDist [5], Cellpose [6] provide pre-trained models e.g. in ImageJ [7-8], QuPath [9], CellProfiler [10] or BIAS [11] plugins convenient for the users, yet in most cases accuracy strongly depends on the image data. When applied to a certain experiment type the pre-trained models do not meet the expectations, and re-training or training from scratch is needed. To this end annotated (labelled) images must be prepared for which available software tools can be used; a recent article reviews them [12]. The challenging part of object annotation is either the number of objects to label or the difficulty of the morphology (shape), texture, border recognition by eye even for experts. To help create annotations, automatic solutions have been added to annotation tools [7-9,12-15] (see also **Fig.1A**). The most challenging task remains 3D object labelling due to the generally low resolution along the z axis in case of z-stack images, the implicitly complicated structure/texture of the sample such as spheroids or organoids (see examples on **Figure 1B-C**) and the large file size of such data.

**Figure 1.**
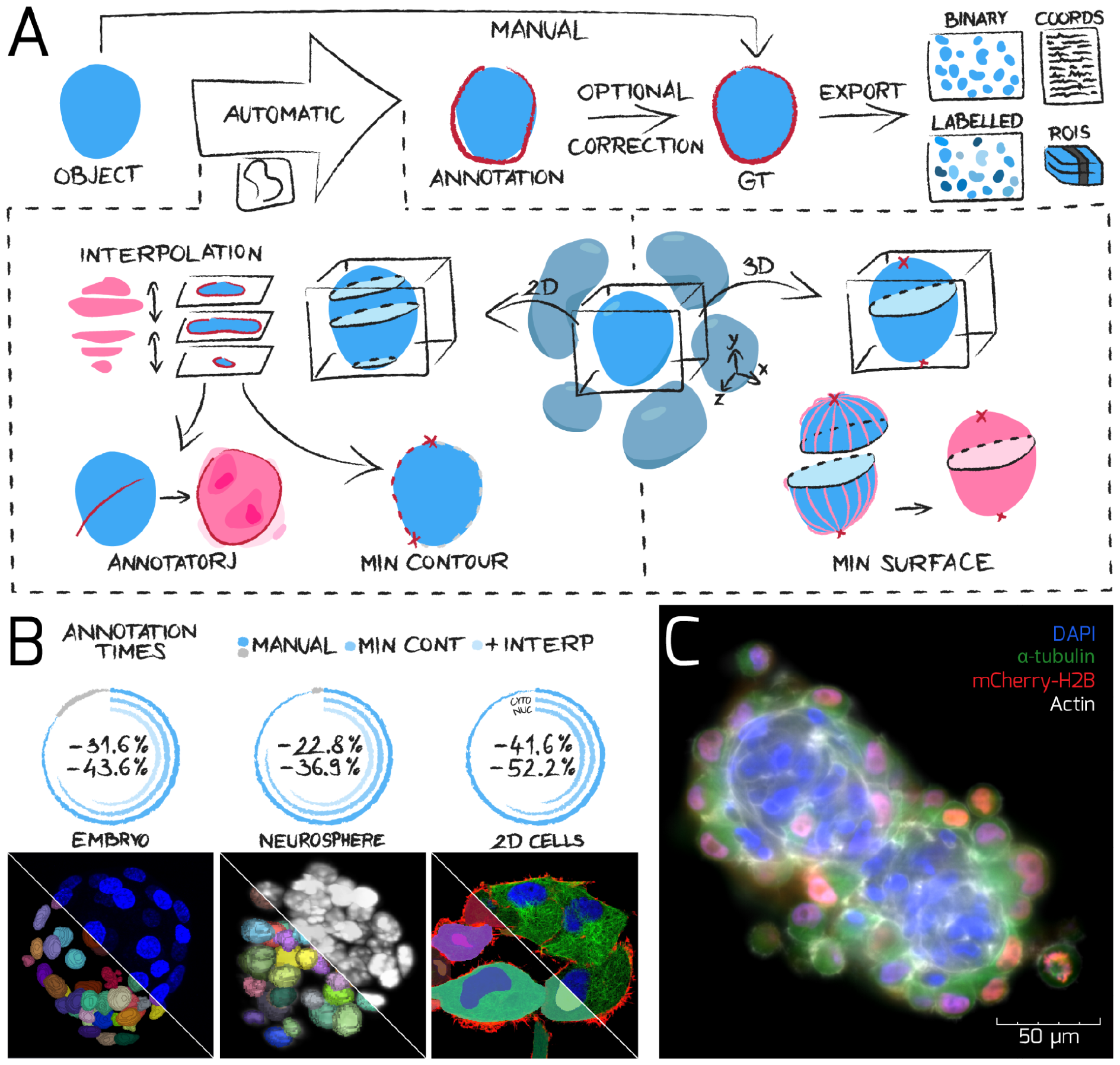
Overview of *napari-biomag-annotator* when labelling single nuclei in a spheroid. A) Schematic representation of the annotation methods. B) Data used for benchmarking the tool with average time saved compared to manual annotation. Datasets are visualized in the napari viewer with annotations overlayed. Pie charts show annotation times using manual, minimal contour or additionally interpolation; gray section corresponds to difference between two sets of manual annotations, while on the 2D cells minimal contour is shown for nuclei and cytoplasm separately. C) Representative co-culture spheroid used in the ground truth dataset at https://doi.org/10.6084/m9.figshare.c.7020531.v1. Abbreviations: GT - ground truth, interp - interpolation.

We present a toolbox of annotation methods implemented in the napari [13] ecosystem entitled *napari-biomag-annotator* as a plugin, freely available at https://github.com/biomag-lab/napari-biomag-annotator and at the napari hub (at https://www.napari-hub.org/plugins/napari-biomag-annotator). Napari supports n-dimensional image inspection in its viewer and several plugins are available for different image processing tasks (e.g. StarDist [5], Cellpose [6] are available as napari plugins). Our toolbox consists of four methods, each aims to assist in single object annotation (presented on **Figure 1**) as follows. 1) *Minimal contour* allows 2D annotation in a few clicks by approximating the contour of the annotated object between two points via image features with high values near object boundaries (e.g. gradient for fluorescent images), 2) *Minimal surface* can be used for 3D annotation similarly to minimal contour by approximating surfaces using as few as two points as reference, 3) *Slice interpolation* is most helpful in z-stack annotation when one object is present on multiple layers and only sparse annotations are available by interpolating contours on unlabelled layers using energy minimization of a functional, 4) *AnnotatorJ* [14] suggests contours around the object using deep learning after an initial contour is quickly drawn. A detailed discussion of the underlying methods is found in **Methods** (section *Annotation tools*) and **Supplementary Note 1** (methods 1-3), see also **Supplementary Videos 1-2**.

The effectiveness of our proposed plugin is demonstrated on microscopy image data arising from six experiments: I) 2D, II) Embryo [16], III) Neurosphere [17], IV) Co-culture, V) Mitotic spheroid data [18] and VI) Melanoma (**Supp. Table 1**). We asked expert annotators with relevant expertise in single-cell labelling on 2D and 3D images to label single cell objects on the datasets using the proposed automatic annotation tools in *napari-biomag-annotator*. The combined dataset contains overall 5861 images with more than 3000 labelled objects including nucleus and cytoplasm annotations. Data I) was used for validation on different cell compartments, while data II) and III) were used as a benchmark where the experts annotated the same objects twice manually to provide an accurate time and intersection over union (IoU) comparison. Datasets IV) and V) were used to provide curated ground truth annotations for 3D images. Dataset VI) served as a test to validate the Python reimplementation of AnnotatorJ. Experimental description of the datasets is provided in **Methods**.

In our experiments, we measured the annotation time and precision according to the IoU metric against manual labelling (*E*) comparing inter- and intra-annotators, as well as against our tools. The usage of *Minimal contour (MC)* by itself and combined with *Slice interpolation (I)* were tested on datasets II-III) and compared against two times repeated manual labelling; the performance of the *Minimal Surface (MS)* tool was assessed on dataset II); *AnnotatorJ* was already benchmarked in an earlier study [14] - thus only a comparison to an existing manual annotation from an expert is shown on dataset VI) (see **Methods**). We have come to the following conclusions (see **Fig. 2**). 1) Manual annotations tend to have an average 0.8825 (II) and 0.8042 (III) intra-expert IoU difference, while 0.8900 (I), 0.8238 (II) and 0.5789 (III) inter-expert (see **Fig. 2A,C,E,G**) on the given datasets matching our expectations based on [14,19,20]. 2) The average precision based on IoU did not change considerably when using either only *MC* (intra-person: I: 0.8870, II: 0.8555, III: 0.7539, inter-person: I: 0.8784, II: 0.8164, III: 0.6153) or together with interpolation (intra-person: II: 0.8481, III: 0.7539, inter-person: II: 0.8079, III: 0.5904) compared to manual labelling (**Fig. 2A,C,E,G, Supp. Fig.1A,D-F**), confirming the reliable performance and usability of the tools when creating ground truth annotations or manual (semi-automatic) segmentation on cell culture images. The *Minimal surface* method was tested only on dataset II), as dataset III) contains small objects which removes the advantage of the technique; yielding an IoU of 0.8032 (**Supp. Fig.1B**). 3) Annotation times were reduced by an average of 22.88% (20.83 minutes per image and 20.35 seconds per object) when using *Minimal contour* compared to manual labelling times (85.76 minutes per image and 94.81 seconds per object). However, when using both tools we could save an additional 26.4% totalling a 49.28% time saved on annotations (42.35 minutes per image and 46.62 seconds per object) (see **Fig. 2B,D,F,H, Supp. Fig.1C**). *Minimal surface* achieved 12.35% (7.97 minutes) acceleration in annotation time compared to the fastest manual time with an extra 10.43 minutes of algorithm runtime. Nevertheless, not every image or annotator resulted in such an ideal time difference: the negative outlier had an 187.92% increase in time on dataset III while the highest annotation time decrease was 85.74% on dataset II. See more detailed results and experiment setup in **Methods** (section *Evaluation*) and **Supp. Fig.1**.

**Figure 2.**
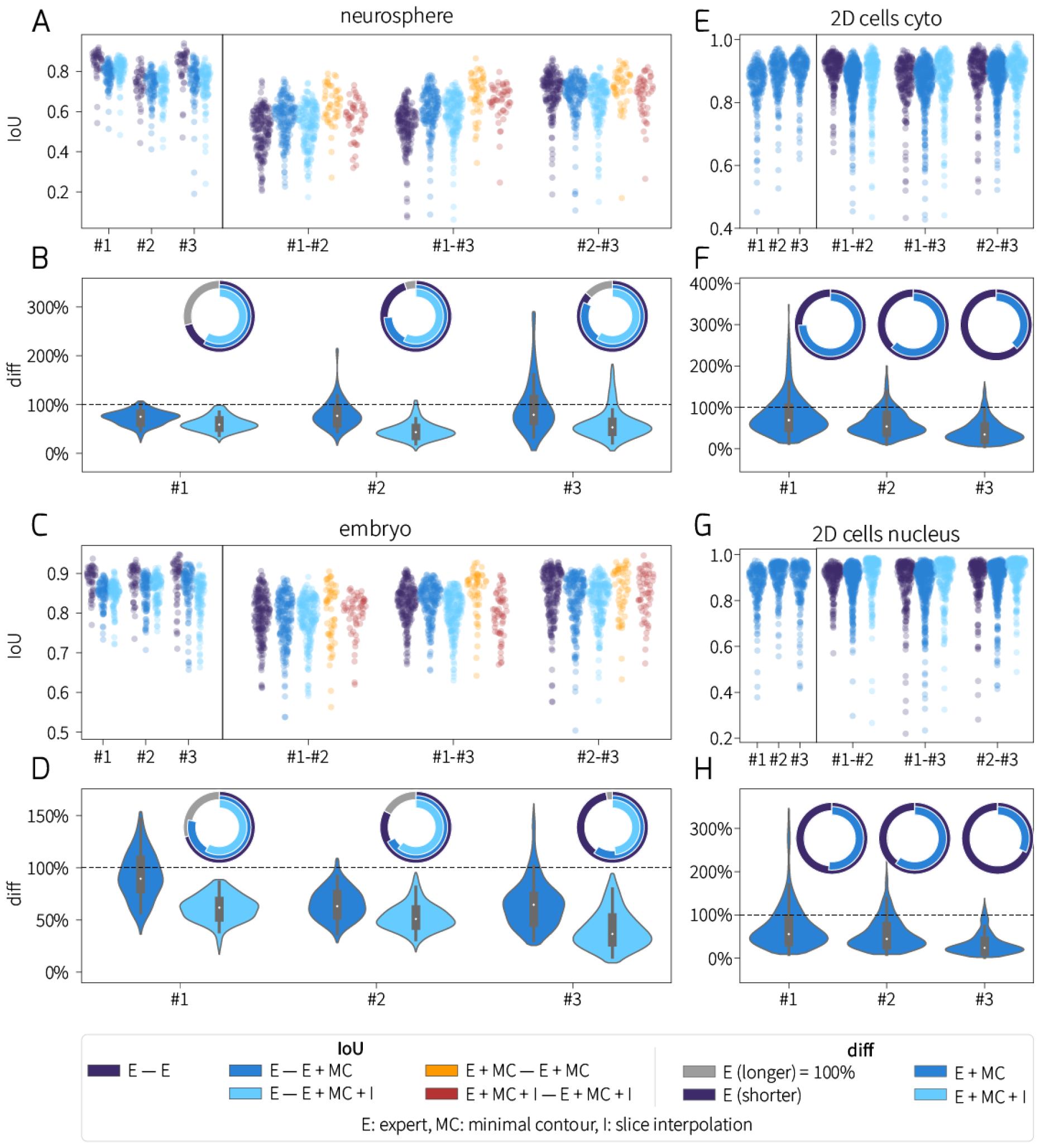
Results on the three benchmark datasets. Swarm-, violin- and pie plots are grouped by datasets as indicated by titles and shown by annotators (experts, #1-3). Comparisons are colour-coded as in the legend, see also abbreviations. A-B) Neurosphere, C-D) embryo, E-F) 2D cells cytoplasm, G-H) 2D cells nucleus data. A,C,E,G) Inter-expert IoU scores were corresponded in a pairwise manner according to the legend in the bottom, insets on the left show intra-expert differences in manual annotation. B,D,F,H) Violin plots (without idle times) and pie charts (total) represent relative annotation time efficiency compared to manual annotation as 100% indicated by dashed line and full circle, respectively. Pie charts show the difference between manual annotations in grey where applicable. See also **Supplementary Fig. 1** for averaged and detailed results on all annotators.

Such a tool is invaluable for the bioimage analyst community as it saves the expert time while making the labelling process more convenient and less exhausting. When training a segmentation model is not feasible (e.g. due to the lack of computational resources), annotations created in *napari-biomag-annotator* can be directly used in downstream analysis such as statistics or further analysis of cellular data. The tools rely on strong mathematical foundations whose efficiency was proven to surpass manual labelling and other popular tools by approximately 38% and 32% in time, respectively (see **Supp. Tables 2-3 and Supp. Fig.1C**) while keeping accuracy on par. The released annotated spheroid dataset may be used for model training in its domain or method development generally.

**Supplementary Figure 1.**
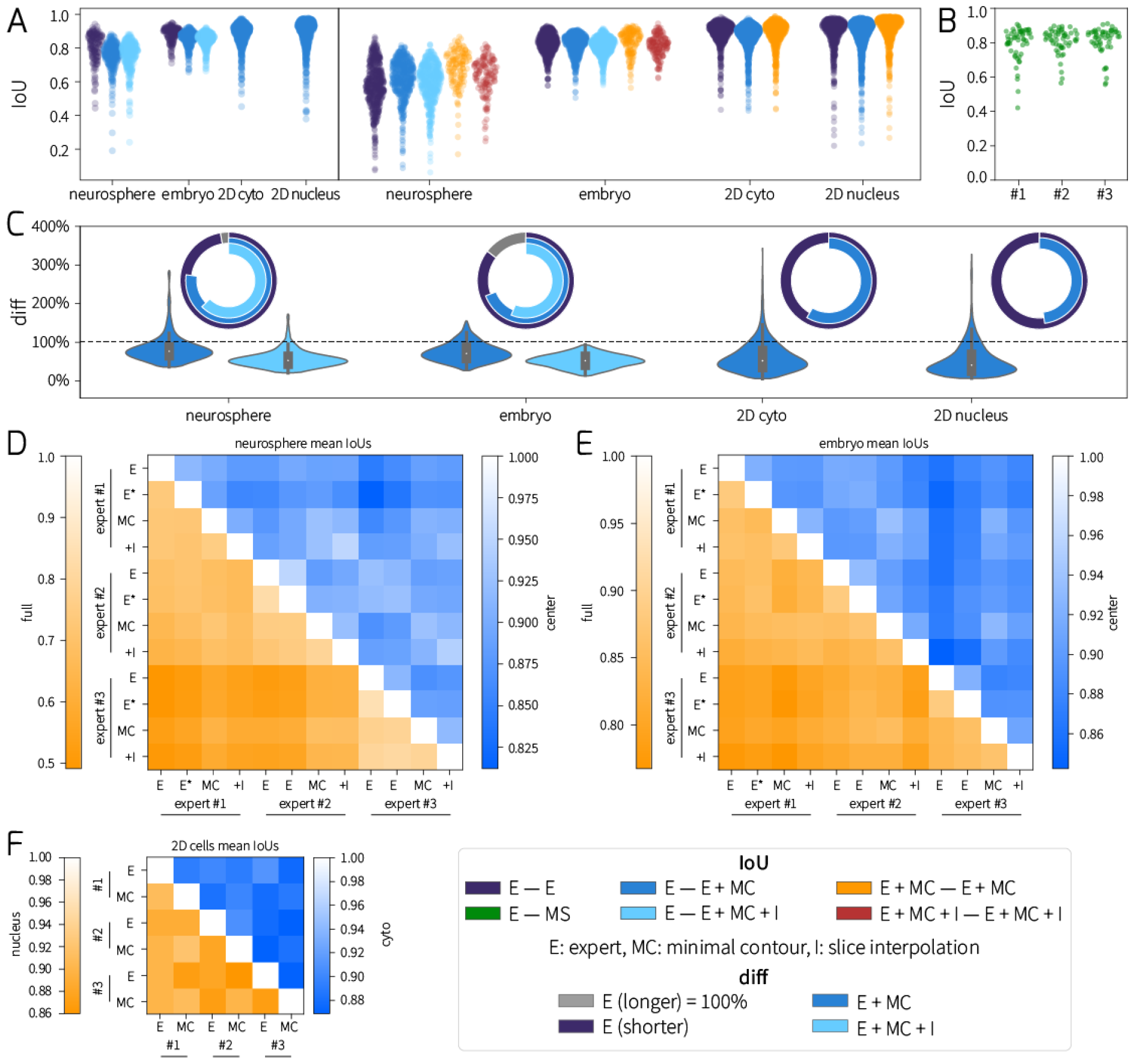
Averaged measurements on datasets and annotators. Visualization and labels are as on **Figure 2** (see also legend in the bottom right). A) Averaged IoU scores plotted in a pairwise manner according to the legend in the bottom of **Fig. 2**, inset on the left shows intra-expert differences in manual annotation. B) Minimal surface measurements. C) Violin plots (without idle times) and pie charts (total) represent relative annotation time efficiency compared to manual annotation as 100% indicated by dashed line and full circle, respectively. Pie charts show the difference between manual annotations in gray where applicable. D-F) Matrices of measured mean IoU scores by annotators and methods, datasets are as on the title and y axis labels next to the colour bars in the upper and lower triangular, respectively; full and center on D-E stand for objects in the entire 3D structure and on the central 50% of slices, respectively, experts are labelled #1-#3 (shortened on F).

## Methods

### Annotation tools

Mathematical foundations of the methods are briefly summarized as follows; for an in-depth discussion see **Supplementary Note 1**.

#### Minimal contour

This tool provides easy and quick annotation in 2 dimensions. Given two or more reference points, our method approximates the optimal curve between these points with variational minimization, thus, annotation of objects with continuous contours can be carried out even with just two clicks. Formally, the task is to find a path between two reference points that minimizes 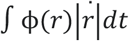, where *r*denotes the curve going between the two reference points and ϕ is a carefully defined function usually denoting image intensity gradient information according to which the optimal curve *r*can be found.

#### Minimal surface

3D surface annotation with as few points as possible is done with the minimal surface algorithm, which is an extension of the minimal contour method. The surface approximation is carried out by optimizing the energy function 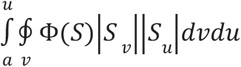, where *S* denotes the surface, however, there is no known solution for this problem as of today, so we reformulate this task to approximate the surface as a set of minimal contours, thus, we optimize multiple curves that lie on the surface in 3D. Technically, two points were applied in the software. The algorithm will also require a 2D annotation of a slice between the two points.

#### Mean contour

We provide a 2-dimensional contour averaging method with this tool which can mainly be used to annotate just a few z-slices while interpolating the ones in-between the annotations. Our method is based on variational optimization, where we represent the annotations as 2D parametric contours. To achieve an optimal approximation between two slices, we find a reparametrization function γ that minimizes 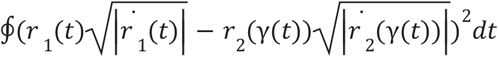, where *r*_1_ (*t*) and *r*_2_ (*t*) denote two annotated slices between which we would like to approximate the contours.

#### AnnotatorJ

Adaptation of the original ImageJ version of AnnotatorJ [14], this tool is intended for 2D object annotation and export offering a convenient contour assist method via prediction with an integrated deep learning model, specifically U-Net [21], based on an initial contour drawn by the user quickly and imprecisely. The initial contour is used to approximate the area where the prediction is desired and returned as a thresholded version of the predicted probabilities. Further functionalities include the training of new models to be used in contour assist, editing of contours, class annotation, import and export to training data formats, annotation types (instance, bounding box, semantic).

## Datasets

In total six image datasets were used to benchmark (datasets I-III) and demonstrate the capabilities and performance of the annotation tools (datasets IV-VI), see **Supp. Table 1** below and https://doi.org/10.6084/m9.figshare.c.7020531.v1.

**Supplementary Table 1.**
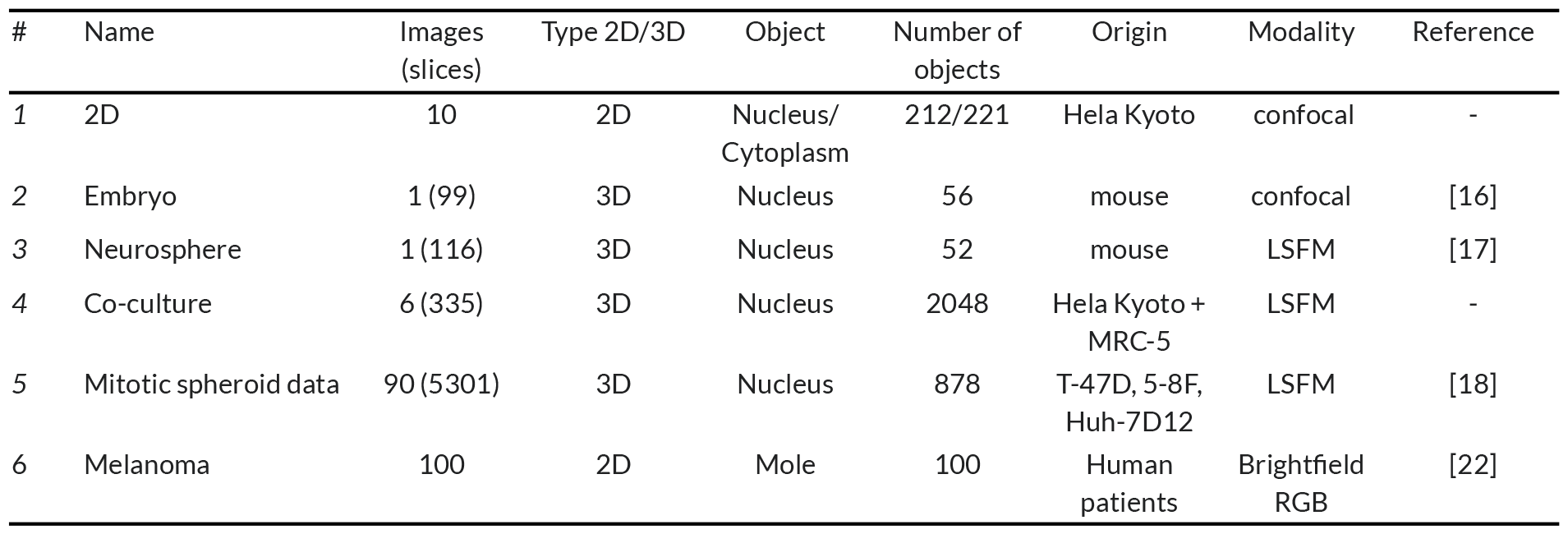
Summary of the datasets used.

### 1. 2D cells

HeLa Kyoto cells were seeded on a glass coverslip and after one day of incubation time the cells were fixed with 4% paraformaldehyde (PFA) and washed with Dulbecco’s Phosphate Buffered Saline (DPBS) and treated with 0.1% TRITON-X for 10 minutes. After that, the cells were washed with DPBS three times, then stained with 1 μg/ml DAPI and 1:200 Flash Phalloidin NIR 647 (424205, Biolegend) dissolved in DPBS for 10 minutes at room temperature. After staining, the cells were washed with DPBS three times and the coverslip was secured on a glass slide for further analysis. For imaging, an Olympus Fluoview FV 1000 microscope was used with a 60x/1.35 objective, and the exposure time and laser power were adjusted for each channel separately (DAPI 405, Alexa Fluor 488, and Alexa Fluor 633. Each fluorescent image is 2048 × 2048 pixel resolution with 0.103 μm pixel size.

This dataset was used as datasets II-III) except only as 2D segmentation training data for nucleus and cytoplasm.

### 2. Embryo data [15-16]

This dataset of a mouse embryo was primarily used for quantitative evaluation of our tools while also creating manual ground truth annotations to be later used as 2/3D nucleus segmentation training data. The dataset contains easily distinguishable objects that offer minimal overlapping regions. Images may be downloaded free of charge from https://www.3d-cell-annotator.org/download.html.

### 3. Neurosphere data [15,17]

This LSFM (light-sheet fluorescence microscopy) dataset was used as dataset II), it contains a 3D image of a small spheroid on 116 slices with nuclei labelled fluorescently. This dataset is valuable since it has a lower resolution with overlapping objects. Images are also free to download from https://www.3d-cell-annotator.org/download.html.

### 4. Co-culture spheroid data

This dataset was not published before.

Three-dimensional co-cultures were generated using the HeLa Kyoto EGFP-alpha-tubulin/H2B-mCherry cervical cancer cells (Cell Lines Service) and MRC-5 fibroblasts (American Type Culture Collection). Cell cultures were maintained following the manufacturer’s instructions. For spheroid generation, we used a co-culture medium consisting of DMEM, 10% FBS (Euroclone), 1% L-glutamine (2mM), and 1% Penicillin-Streptomycin-Amphotericin B mixture (all from Lonza). First, 60 cancer cells were seeded into each well on U-bottom, cell-repellent 384-well plates (Greiner Bio-One); and after 24 hours of incubation at 37 °C and 5% CO_2_, 240 fibroblasts per well were added onto the HeLa cells. After 24 hours, the co-culture spheroids were collected, washed three times with Dulbecco’s Phosphate Buffered Saline (DPBS), fixed with 4% PFA for 60 min, then washed again with DPBS three times, and stored at 4°C in DPBS until imaging. Before imaging, spheroids were incubated in 1% Triton X-100 overnight at room temperature and washed three times with DPBS. For labelling, spheroids were stained with 1 μg/ml DAPI overnight, then 1:200 Flash Phalloidin NIR 647 (424205, Biolegend) was applied for 60 min. In the end, spheroids were washed with DPBS three times before imaging. The preparation for imaging and all the imaging parameters were the same as we discussed in the data article [18]. The Leica SP8 Digital LightSheet microscope was used to create fluorescent images of each spheroid. The images were taken with 200 ms exposure time with adjusted laser intensity for each channel at 405, 488, 552, and 638 nm (maximum laser intensity 350 mW), and a 25x/0.95 detection objective was used for the light-sheet imaging with the 2.5 mm mirror device on the objective. For each spheroid, dH_2_O mounting medium was used. The images have a 2048 × 2048 pixel resolution with 0.14370117 μm pixel size and with a 3.7 μm distance between the images in each z-stack. This dataset is intended to be used for 2/3D nucleus segmentation training purposes, to specifically target 3D spheroids. To decrease the blurry effect of light scattering inside of the co-culture spheroids, LIGHTNING was used as a post-processing step (available with the LAS-X 4.4 software, Leica).

### 5. Mitotic spheroid data [18]

This dataset was used to create ground truth annotations for the particularly problematic mitotic nucleus phenotype which tends to cause problems for automatic segmentation methods due to the intrinsically complicated geometry of condensed DNA. The dataset includes 90 multicellular cancer spheroids derived from 3 cell lines (i.e. T-47D, 5-8F, and Huh-7D12) with a diameter of 250±30 μm. The images have 1 channel for the fluorescently (DRAQ5-ThermoFisher, USA) labelled nucleus and were acquired with a light-sheet microscope. This dataset is intended to be used for 2/3D nucleus segmentation training purposes, to specifically target dividing cells. Overall 878 objects were annotated and classified as dividing cells. Images are also free to download from https://doi.org/10.6084/m9.figshare.12620078.v1.

### 6. Melanoma [22-23]

The HAM10000 dataset consists of high-resolution 2D colour images from different populations in RGB format, each with metadata containing the patients’ previous health records. The images are taken by dermatologists using a dermoscope and capture different skin lesions from different patients, covering all major diagnostic categories related to pigmented moles. Each image represents a distinct skin lesion, accurately labelled with the corresponding dermatological modification (benign or malignant), validated by dermatologists. For cases without histopathological confirmation, ground truth class label was established by follow-up, expert consensus or confirmation by in vivo confocal microscopy. The entire dataset can be downloaded from https://doi.org/10.7910/DVN/DBW86T and used for academic purposes.

A subset of 100 images was selected to include both benign and malignant cases. These images were annotated by outlining the lesion and separating the object under examination from the background using our annotation tool. This dataset can be used as segmentation training data for example in mAIskin, an automated melanoma detection system that supports and simplifies the work of dermatologists.

## Experiments

### Annotation strategy

Datasets I-III) were used to evaluate the effectiveness and time consumption of our assisted annotation napari plugin tools compared to manual annotations. Three field experts with relevant background and experience in cell and nucleus identification on microscopy images were asked to do the experiments as follows. Each participant labelled every object on every image or slice of a 3D image such that the experts discussed and agreed upon a common annotation strategy regarding which image object to identify as a single nucleus or touching adjacent nuclei, especially in mitotic cases, and the approximate brightness and contrast settings in napari so that the annotations could be comparable and the created ground truth annotated datasets (datasets I, IV-V) could be consistently used for training purposes. Still, differences occurred even intra-expert when the same person annotated the images twice due to the natural effect of tiring of the human eye and decreased focus and patience completing a repetitive and long task; also see **Results**.

The annotators labelled all images manually twice; this was the basis of our comparative measurements of both time and accuracy intra- and inter-expert and against the tools. Then, experts labelled the images using 1) only *Minimal contour*, 2) together with *Slice interpolation. AnnotatorJ* has already been quantified in [14] earlier thus we did not repeat the same experiment on its napari plugin.

When using only *Minimal contour* the experts utilized the image feature-based edge detection capability of the tool, having only to place a few points on the object contour for the tool to extend the path between the points creating a closed curve around the border of the object.

*Slice interpolation* allowed several 3D slices to be skipped when annotating the same object extending to multiple z-stack slices and the contour on the missing slices interpolated. This method especially reduces annotation time when the resolution in z is high i.e. an object is present on a high number of z slices. The most efficient annotation strategy was using both tools.

For the embryo data, the following parameters were used: Param: 6, Blur sigma: 0.2, Smooth contour: 0.75, while for the neurosphere data Param: 8, Smooth contour: 0.52.

### Quantitative evaluation of annotation tools

Experiments were quantitatively evaluated using the classical definition of IoU (intersection over union) determining the ratio of pixels corresponding to the overlapping area between the two objects i.e. intersection and their union including all pixels of the two objects. This pixel-level calculation of quality assessment results in a single floating point-precision score for annotation pairs of the same object while object-level evaluation (see TS referred as mAP_2_ in [24] below) returns the counts of TP (true positive), FP (false positive) and FN (false negative) objects on the image then calculates a single IoU score for the image based on an overlap threshold being considered as a positive detection. Using the following pixel-level metric for IoU

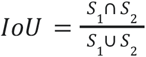

where *S*_1_, *S*_2_ are the set of pixels marked as object pixels by two annotators, accuracy scores are yielded for all objects which then can be aggregated to e.g. averaged IoU scores such as those displayed on **Supp. Fig. 1D-F**.

Additionally, annotations were also assessed using the following definition of threat score (TS) which shows the ratio of TP, FP and FN objects commonly applied in standard computer vision problems such as object detection or instance segmentation. We rely on the following definition of threat score formulated as follows

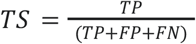

where TP is the number of labelled objects with an overlap to the ground truth object above a given threshold, FP is the number of labelled objects that have no corresponding ground truth object with overlap above the given threshold and FN is the number of ground truth objects that have no corresponding labelled object with overlap above the given threshold. We used thresholds between 0.5 and 0.95 with steps of 0.05 in our evaluation.

Each annotation was compared to all other annotations one by one resulting in a matrix of average IoU scores. Annotation times were similarly compared. Results are represented as heatmaps on **Supp. Fig. 1D-F**.

Annotation times were measured inside the plugin on object level considering long pauses above 10 seconds as idle time such that only relevant times are quantified when the expert is creating an annotation for an object. These filtered times are displayed as violin plots on **Fig.2B,D,F,H** and **Supp. Fig.1C**, while the total time taken to annotate all objects on the entire image from start to finish including idle times is displayed on pie charts (see also **Fig.1B**).

One of the challenges of manual annotation, outlining complex borders, can be observed in dataset I). Cells adhere to the surface of the culture container therefore their borders (visualized by the Phalloidin channel) become irregular and difficult to manually track. Thus, assisted annotation is especially advantageous in single-cell cytoplasm contouring.

### Ground truth data

As the fundamental purpose of the annotation tool is to create ground truth annotations that can later be used for training, we demonstrated this capability by labelling datasets IV-V) made freely available at https://doi.org/10.6084/m9.figshare.c.7020531.v1. Dataset V) of the mitotic nucleus annotations fills a niche in open annotated datasets in its domain. Whereas dataset IV) presents co-culture spheroid annotations to which similar open annotated dataset the authors have not found as of writing. Both datasets can be easily used to train new single-cell segmentation models or start developing new methods.

## Results

### Annotation tool comparison

In previous studies [12,14,25] we have conducted comparative analysis of annotation software available to the target community focusing on free and open source tools. In addition to those detailed before, further tools are shown below in **Supp. Table 2** and detailed as follows.

**Supplementary Table 2.**
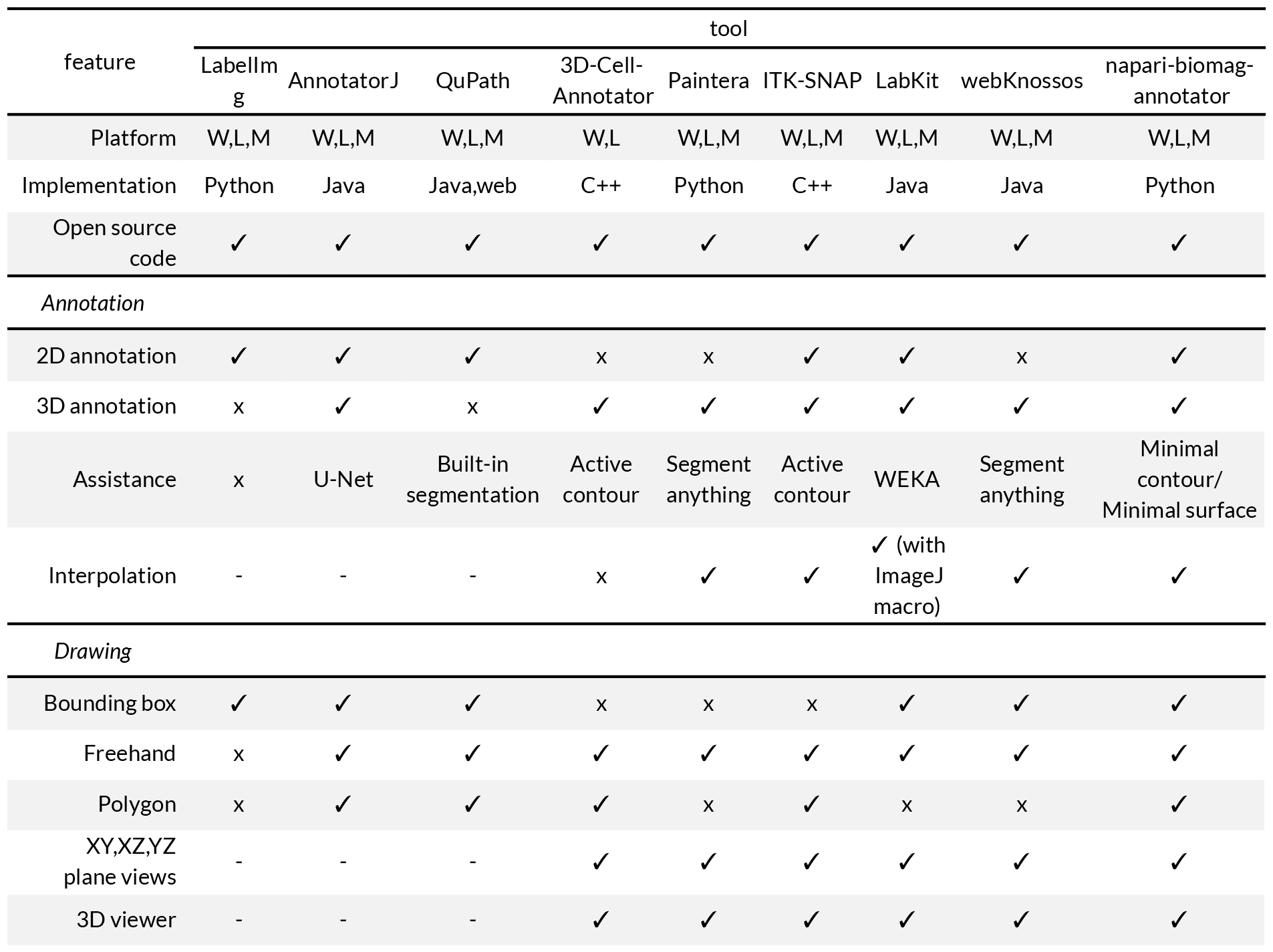
Comparison of annotation software tools.

| feature | tool |  |  |  |  |  |  |  |  |
| --- | --- | --- | --- | --- | --- | --- | --- | --- | --- |
|  | Labellm<br>g | AnnotatorJ | QuPath | 3D-Cell-Annotator | Paintera | ITK-SNAP | LabKit | webKnossos | napari-biomag-annotator |
| Platform | W,L,M | W,L,M | W,L,M | W,L | W,L,M | W,L,M | W,L,M | W,L,M | W,L,M |
| Implementation | Python | Java | Java,web | C++ | Python | C++ | Java | Java | Python |
| Open source<br>code | ✓ | ✓ | ✓ | ✓ | ✓ | ✓ | ✓ | ✓ | ✓ |
| *Annotation* |  |  |  |  |  |  |  |  |  |
| 2D annotation | ✓ | ✓ | ✓ | x | x | ✓ | ✓ | x | ✓ |
| 3D annotation | x | ✓ | x | ✓ | ✓ | ✓ | ✓ | ✓ | ✓ |
| Assistance | x | U-Net | Built-in<br>segmentation | Active<br>contour | Segment<br>anything | Active<br>contour | WEKA | Segment<br>anything | Minimal<br>contour/<br>Minimal surface |
| Interpolation | - | - | - | x | ✓ | ✓ | ✓ (with<br>ImageJ<br>macro) | ✓ | ✓ |
| *Drawing* |  |  |  |  |  |  |  |  |  |
| Bounding box | ✓ | ✓ | ✓ | x | x | x | ✓ | ✓ | ✓ |
| Freehand | x | ✓ | ✓ | ✓ | ✓ | ✓ | ✓ | ✓ | ✓ |
| Polygon | x | ✓ | ✓ | ✓ | x | ✓ | x | x | ✓ |
| XY,XZ,YZ<br>plane views | - | - | - | ✓ | ✓ | ✓ | ✓ | ✓ | ✓ |
| 3D viewer | - | - | - | ✓ | ✓ | ✓ | ✓ | ✓ | ✓ |

Recently an in-browser annotation tool was published for 3D electron microscopic data called webKnossos [26]. For comparison, webKnossos was tested on the embryo dataset (data II) where accuracy and time were measured. Based on the feedback of the expert, webKnossos is a user-friendly tool that requires no parametrization. Exceptional performance was noted with blurry and less visible objects, while the automatic detection utilizing the implemented AI model was less convincing in the case of dividing/mitotic cells. The AI-based prediction frequently connects small, nearby objects that are not in contact, necessitating human correction. Furthermore, a single object prediction often takes more than 1 or 2 seconds (**Supplementary Video 2**) which greatly increases the annotation time. To enhance the performance, an interpolation method is available, however, it can only interpolate between the last two contours.

Considering that webKnossos was developed for 3D electron microscopic data and more models are under development, it offers an easy-to-use solution for annotation.

ITK-SNAP (Insight Segmentation and Registration Toolkit for Semi-Automatic Segmentation of Structures in Medical Images) is an open-source software application that is a part of the broader ITK initiative, which aims to provide a collection of software tools for medical image analysis and computational anatomy [27]. In order to compare this tool to our proposed one, one of our experts annotated the embryo dataset with ITK-SNAP (**Supplementary Video 2**). The expert acknowledged the utility of ITK-SNAP as a valuable tool. However, ITK-SNAP may have a steeper learning curve, especially for users who are new to medical image analysis or 3D annotation tools. The software offers a variety of features, and users may need some time to become familiar with its functionalities. During annotation the expert found that the active contour method (that is employed within the tool as part of a region growing process) works with high precision even in blurry parts of the original image.

Paintera (https://github.com/saalfeldlab/paintera) is a visualization and annotation tool that was developed to handle large-scale volumetric data, such as those generated by various imaging modalities in the field of connectomics. Again, we used the embryo dataset as a base to compare this tool to *napari-biomag-annotator*. The assessment of the expert indicates that this tool has many useful annotation functionalities such as live visualization of interpolation between slides or built-in keyboard shortcuts. However, there is a limitation of this tool, namely it can import only N5, HDF7, and Zarr files and can export only in its own data format.

For an in-depth comparison our experts annotated dataset II) with the aforementioned tools: expert #1 with Paintera and ITK-SNAP, and expert #2 with webKnossos (**Supp. Table 3**). We found that using Paintera we got a similar precision compared to our semi-automatic methods, but the annotation time remained the same. As for ITK-SNAP we have seen a small drop in the IoU score, the annotation time was significantly higher (∼+40%) than manually. Only webKnossos was able to achieve similar annotation time as our *Minimal contour* method but was slower than *Minimal contour* with *interpolation* while the IoU score was inferior to ours.

**Supplementary Table 3.**
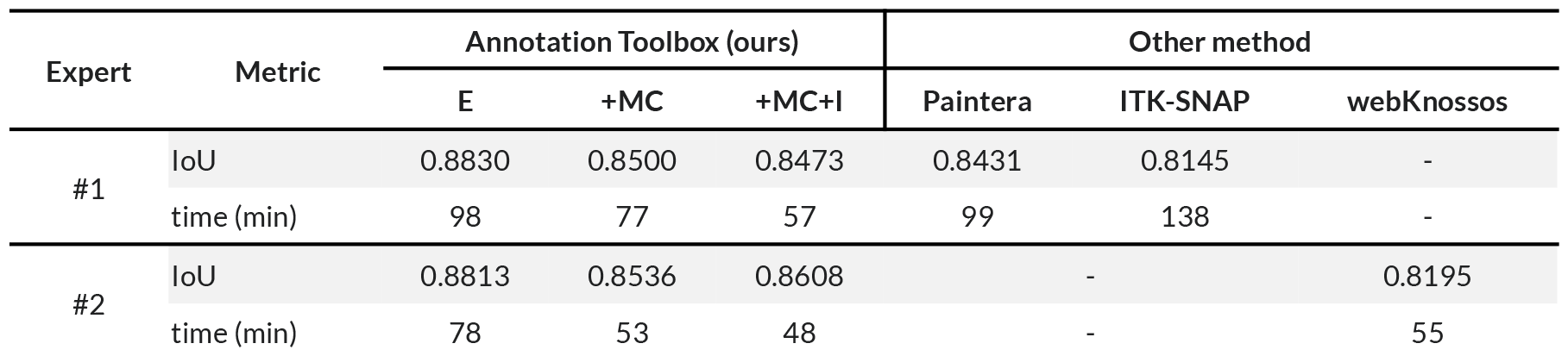
Assessment of other annotation tools compared to ours.

### Evaluation

Annotation accuracies according to the classical definition of IoU and times are represented on **Supplementary Figure 1A** and **C**, respectively. IoU scores were aggregated for all three expert annotators on **Supp. Fig. 1A** and are displayed by datasets whereas on **Fig. 2A,C,E,G** scores are compared between the experts (inter-expert). The matrices in **Supp. Fig. 1D-F** represent individual IoU scores for the three experts and the applied methods, also comparing the entire stack of 3D images in case of datasets II-III) (denoted as full) and the central 50% of slices in the z-stack (center) corresponding to the given object. Obviously, scores are generally higher in the central region for both datasets.

For intra-person similarity (i.e. the IoU score between different annotations of the same annotator) we experienced a relative difference of II) -3.06% and III) -6.06% using minimal contour, while it was II) -3.89% and III) -6.10% when combining *Minimal contour* and *Slice interpolation*. 2) The inter-person similarity (that is the IoU score between different annotators) was I) -2.09%, II) -0.79% and III) +8.78% with *Minimal contour*, while with *Minimal contour* and *interpolation* II) -1.76% and III) +4.27% was achieved. Using the *Minimal surface* method the inter-person IoU score change was on average II) -2,42%, without any manual correction.

*Minimal surface* was tested on dataset II) since the method is intended for larger 3D objects while dataset III) comprised small nuclei and our third dataset used for benchmarking (dataset I) was 2D, concluding the annotation accuracies remained comparable to manual labelling, thus proving the efficiency of the method. See **Supp. Fig 1.B**. Additionally, the time required by the annotator operating the plugin with the *Minimal surface* method decreased by 12.35%.

We inspected the most extreme outliers in our dataset, both with the highest deterioration and the largest improvement in regards to annotation time. The negative outlier was an 187.92% increase in annotation time on dataset III). For this object the annotation time using *Minimal contour* was near average, while manually it was the fastest annotated object with one third of the average annotation time. This was a rather difficult object as it was both blurry and touching another object which is reflected in the IoU as well: the intra-person IoU was 0.5122 (0.4708 for the annotator in question), and the inter-person IoU was 0.2134 for manual annotation. On the other hand, the highest annotation time decrease was 85.74% on dataset II) using *Minimal contour* and *interpolation*. This was an easy task for the *Minimal contour* method, as the object had clear contours, thus the IoU was not affected: 0.8592 and 0.8930 against the two manual annotations of the same annotator (compared to 0.8918 between the two manual annotations). Note this annotator had less experience in image annotation which shows the power of the method guiding annotators new to the task.

Additionally, we conducted evaluation of the performance of *AnnotatorJ* on dataset VI) representing a non-microscopy image domain, yielding an average IoU of 0.9700 with thresholds in [0.5-0.8] and 0.7820 with thresholds in [0.5-0.95] by steps of 0.05 according to the object-level definition of IoU (referred as TS or mAP_2_ in [24]). Annotation times were reduced by an average 77.94% (3.63 seconds per object from 16.46), aligned with our expectations from a previous study [14]. As a comparison, using the same object-level assessment on dataset II), the average IoU with *Minimal contour* and *interpolation* was 0.8333 and 0.6575 with thresholds in [0.5-0.8] and [0.5-0.95], respectively, while without interpolation 0.7774 and 0.6176. Manual labelling yielded 0.8214 and 0.6814 on the above threshold ranges.

## Acknowledgments

We acknowledge support from the Lendület BIOMAG grant (no. 2018–342), TKP2021-EGA09, Horizon-BIALYMPH, Horizon-SYMMETRY, Horizon-SWEEPICS, CZI Deep Visual Proteomics, H2020-Fair-CHARM, HAS-NAP3, the ELKH-Excellence grant from OTKA-SNN no. 139455/ARRS, the FIMM High Content Imaging and Analysis Unit (FIMM-HCA; HiLIFE-HELMI), and Finnish Cancer Society. RH and DB acknowledge help from CZI’s “napari Plugin Foundations Grants (Cycle 1 and 2)”. The data management and uploading to FigShare were aided by Filippo Piccinini, for which the authors are grateful. The authors would like to thank Judith Reddington for providing the LIGHTNING processed co-culture images. Authors are thankful to Michael Barbier and Winnok De Vos for their help in developing the Slice interpolation method.

