## Supplementary Note 1 for "When the pen is mightier than the sword: semi-automatic 2 and 3D image labelling"

### 1 Mathematical Background

The following mathematical discussion is József Molnár's intellectual property. We cite his articles on the Minimal Surface Problem [2] and the Mean Contour Problem [1]. The reader is encouraged to read his articles in unison to aid the understanding of the following Sections.

#### 1.1 Vector Calculus and Variational Calculus

Vector calculus concerns itself with the differentiation and integration of scalar fields and vector fields. This is primarily done in the 3-dimensional Euclidean space, but we shall state the relevant lemmas and definitions in greater generality, in the $n$-dimensional Euclidean space $\mathbb{R}^n$. Image segmentation requires the definition of parametric curves, so let us first define what we mean by them mathematically.

**Definition 1.1** (Parametric curve)**.** *The points of a parametric curve or path in $\mathbb{R}^n$ are identified with the collections of the differentiable co-ordinate functions of the bounded parameter $\tau$:*

$$q_1(\tau),\ q_2(\tau),\ \ldots,\ q_n(\tau), \qquad \textit{where} \qquad t_1 \leq \tau \leq t_2.$$

The co-ordinate functions may be the well-known Cartesian co-ordinates, but if our problem possesses a certain symmetry, other co-ordinates may be advantageous, e.g. polar or spherical co-ordinates. The rate of change of the co-ordinate functions is described by their *velocity.*

**Definition 1.2** (Velocity)**.** *The velocity of a parametric curve is identified with the derivatives of the co-ordinate functions:*

$$\dot{q}_1(\tau) := \frac{\mathrm{d}q_1}{\mathrm{d}\tau},\ \dot{q}_2(\tau) := \frac{\mathrm{d}q_2}{\mathrm{d}\tau},\ \ldots,\ \dot{q}_n(\tau) := \frac{\mathrm{d}q_n}{\mathrm{d}\tau}.$$

Note that this definition of velocity does not coincide with the velocity definition from classical mechanics. Those familiar with analytical mechanics may recognize this formulation of the parametric curve and its velocity as the generalized co-ordinates and generalized velocity. For the sake of brevity, we assume this generalized framework without mentioning it explicitly.

So far we have been discussing co-ordinates and co-ordinate functions, but we have not mentioned the 'real' vectors of $\mathbb{R}^n$ which will be denoted by boldface symbols. In particular, the position vectors are denoted by $\boldsymbol{r}(\tau)$ and they form a Hilbert space equipped with a norm induced by the dot product. We set up the correspondence between the co-ordinates and the vectors as

$$\boldsymbol{r}(\tau) = \boldsymbol{r}\left(q_1(\tau),\ q_2(\tau),\ \ldots,\ q_n(\tau),\ \tau\right) \qquad \Rightarrow \qquad \dot{\boldsymbol{r}}(\tau) = \frac{\mathrm{d}\boldsymbol{r}}{\mathrm{d}\tau} = \sum_{i=1}^{n} \frac{\partial \boldsymbol{r}}{\partial q_i} \cdot \dot{q}_i(\tau) + \frac{\partial \boldsymbol{r}}{\partial \tau},$$

using the chain rule and abusing the notation. Hence the co-ordinates at particular parameter values can be identified with the vectors themselves, and so can the velocities with the total rate of change of the position vector. This very notion will be helpful in defining the *scalar-by-vector derivative.*

**Definition 1.3** (Scalar-by-vector derivative)**.** *Given a scalar field $U$, its derivative with respect to the co-ordinate $q_i$ is denoted by $\partial U/\partial q_i$. If the co-ordinates are Cartesian, then the vector derivative in the Cartesian basis is*

$$\frac{\partial U}{\partial \boldsymbol{r}} = \sum_{i=1}^{n} \boldsymbol{e}_i \frac{\partial U}{\partial x_i}.$$

*If the co-ordinates are non-Cartesian, then a change of basis linear map transforms the derivative into the appropriate co-ordinate system.*

If a scalar field has no parameter dependence, then its derivative with respect to the position vector is also its total rate of change. If the scalar field does have parameter dependence, then its total rate of change is given by its *gradient*.

**Definition 1.4** (Gradient)**.** *Denoting the Euclidean dot product with $\cdot$, the gradient $\boldsymbol{\nabla} U$ of the scalar field $U$ is related to its total differential:*

$$\mathrm{d}U = \boldsymbol{\nabla} U(\boldsymbol{r}) \cdot \mathrm{d}\boldsymbol{r}.$$

*For this reason, we may occasionally write $\mathrm{d}U/\mathrm{d}\boldsymbol{r}$ for the gradient. This will be based on whether we want to emphasize the vector dependent or independent nature of the gradient.*

An important corollary is that $\boldsymbol{\nabla} U$ is perpendicular to the lines $U = \text{const.}$, called the level sets of $U$. To us, the rate of change of the vector norm will be of great importance, hence it deserves its own Lemma.

**Lemma 1.** *Since the Euclidean norm $\|\cdot\|$ is defined via the dot product, its derivative is*

$$\frac{\partial \|\boldsymbol{r}\|}{\partial \boldsymbol{r}} = \frac{\boldsymbol{r}}{\|\boldsymbol{r}\|},$$

*and similarly to all other 'real' vectors of $\mathbb{R}^n$.*

*Proof.* Write

$$\frac{\partial \|\boldsymbol{r}\|}{\partial q_i} = \frac{\boldsymbol{r}}{\|\boldsymbol{r}\|} \cdot \frac{\partial \boldsymbol{r}}{\partial q_i} = \frac{\boldsymbol{r}}{\|\boldsymbol{r}\|} \cdot \frac{\partial \dot{\boldsymbol{r}}}{\partial \dot{q}_i},$$

and the result follows after substituting in the Cartesian co-ordinates. □

We are now ready to discuss variational calculus, starting with the very definition of *functionals*.

**Definition 1.5** (Functional)**.** *A functional is a mapping that assigns a real number to a function. One such functional $\mathcal{F}$ is of type*

$$\mathcal{F}[q_i] = \int_{t_1}^{t_2} f\left(q_i(\tau),\, \dot{q}_i(\tau),\, \tau\right) \mathrm{d}\tau, \tag{1}$$

*where $f$ is a function and the endpoints of the integral are fixed. The functional depends on all the co-ordinates $q_i$, but only the $i$-th one is written for brevity.*

Functionals are denoted by calligraphic letters and their arguments are enclosed in square brackets by convention. Numerous types of functionals can be defined over function spaces, but we shall restrict our discussion in this section for brevity: hereinafter, by functionals, we only refer to functionals of type (1), even if not explicitly stated. The change in functionals is tracked by the *functional differential.*

**Definition 1.6** (Functional differential)**.** *Given a functional of type* (1)*, its functional differential or variation in the direction of $h$ is denoted by $\delta\mathcal{F}$, and*

$$\delta\mathcal{F}[q_i, h] := \int_{t_1}^{t_2} f\left(q_i(\tau) + \varepsilon h(\tau),\, \dot{q}_i(\tau) + \varepsilon \dot{h}(\tau),\, \tau\right) - f\left(q_i(\tau),\, \dot{q}_i(\tau),\, \tau\right) \mathrm{d}\tau,$$

*where $\varepsilon$ is a small parameter. For small enough $\varepsilon$, the function $f$ can be Taylor expanded.*

#### 1.2 The Quantities and Equations of Motion

Solving problems with variational methods in Computer Science requires the definition of a scalar function called the *Lagrangian*, denoted by $L$. It is a function of the co-ordinates, their derivatives and the parameter:

$$L := L(q_i, \dot{q}_i, \tau).$$

Hereinafter, the dependence on the co-ordinates and their derivatives will be denoted by only the $i$-th co-ordinate and its derivative. The fixed endpoint integral of the Lagrangian is called the action, denoted by $\mathcal{A}$, and it is a functional of $q_i$, the path taken:

$$\mathcal{A}[q_i] := \int_{t_1}^{t_2} L(q_i, \dot{q}_i, \tau)\, \mathrm{d}\tau.$$

Different problems require different Lagrangians and thus yield different actions. If the problem at hand is image segmentation, then the Lagrangian will depend on the image co-ordinates and the image intensity or its gradient values. The true path of a system with action $\mathcal{A}$ is based on the *Principle of stationary action.*

**Theorem 1** (Principle of stationary action)**.** *The path $\xi$ between the fixed endpoints is such that the functional differential of the action $\mathcal{A}$ vanishes at $\xi$:*

$$\delta\mathcal{A}[\xi] = 0.$$

*Such paths are called extremal paths. Since the extremum is almost always a minimum, they are commonly called minimal paths.*

Instead of tackling functional differentials directly, we instead resort to the equivalent Euler-Lagrange Equations which contain ordinary and partial derivatives and connect the problem directly to the Lagrangian. The derivation of the Euler-Lagrange Equations is omitted in this section because it will be discussed in Section 3 at length.

**Theorem 2** (Euler-Lagrange Equations)**.** *The extremal path with co-ordinates $q_i$ between the fixed endpoints is such that its Lagrangian $L$ satisfies*

$$\frac{\partial L}{\partial q_i} - \frac{\mathrm{d}}{\mathrm{d}\tau}\left(\frac{\partial L}{\partial \dot{q}_i}\right) = 0.$$

The Euler-Lagrange Equations are powerful tools for describing systems of many kinds, not only in Computer Science, but also in physical and biological phenomena. However, they are only applicable to systems with fixed initial and end states. For variable endpoint systems, the so-called *Hamilton-Jacobi* framework is used, which is based on a dual formulation, starting with the Hamiltonian

$$H\left(q_i, \frac{\partial L}{\partial \dot{q}_i}, \tau\right) := \sum_{i=1}^{n} \frac{\partial L}{\partial \dot{q}_i} \cdot \dot{q}_i - L,$$

the Legendre transform of the Lagrangian. The quantity $\partial L/\partial \dot{q}_i$ is called the $i$-th *generalized momentum* co-ordinate and it is usual to denote it with $p_i$. However, we refrain from this abbreviation for the reason that we would like to stress its Lagrangian origin. With the introduction of the Hamiltonian, the Euler-Lagrange Equations decouple into three sets of equations called *Hamilton's equations.*

**Theorem 3** (Hamilton's Equations)**.** *The fixed endpoint formulation in the Hamiltonian point of view is:*

$$\frac{\partial H}{\partial q_i} = -\frac{\mathrm{d}}{\mathrm{d}\tau}\left(\frac{\partial L}{\partial \dot{q}_i}\right), \qquad \frac{\partial H}{\partial(\,\partial L/\partial \dot{q}_i\,)} = \dot{q}_i, \qquad \frac{\partial H}{\partial \tau} = -\frac{\partial L}{\partial \tau}.$$

Hamilton's equations can be derived from the Euler-Lagrange Equations and they provide a different aspect to the same problem: these equations may be easier to solve than the original Euler-Lagrange Equations. Now we are ready to take the intellectual leap of extending our framework to variable endpoint systems.

**Definition 1.7** (Hamilton's principal function)**.** *The Hamilton's principal function, denoted by $S$, is a variable upper endpoint integral whose values are obtained when the action is evaluated at the extremal paths $\xi$ between the endpoints:*

$$S(q_i(t), t) := \mathcal{A}[\xi] = \int_a^t L(\xi,\, \dot{\xi},\, \tau)\, \mathrm{d}\tau.$$

The lower bound is an arbitrary constant. Like most authors, we will not take great care in verbally distinguishing the action from the Hamilton's principal function: the reader should be aware of the differences based on the definition above and the context. We are now fully equipped to write down the true evolution equations of a system with variable endpoints.

**Theorem 4** (Hamilton-Jacobi Equations)**.** *The derivatives of the action are related to the Lagrangian and the Hamiltonian:*

$$\frac{\partial S}{\partial q_i} = \frac{\partial L}{\partial \dot{q}_i}, \qquad \frac{\partial S}{\partial t} = -H. \tag{2}$$

The proof of the Hamilton-Jacobi Equations can be found in any standard textbook on variational methods, based on the Euler-Lagrange Equations and Hamilton's Equations.

#### 1.3 Geometric Actions

Geometric actions do not depend on the parameterization of the contour, only on the image content and the intrinsic geometric properties of the contour. An action $\mathcal{A}$ is geometric, if it is invariant under reparameterizations:

$$\int_{t_1}^{t_2} L(q_i(\tau),\, \dot{q}_i(\tau),\, \tau)\, \mathrm{d}\tau = \int_{\gamma(t_1)}^{\gamma(t_2)} L\big(q_i(\gamma),\, \dot{q}_i(\gamma)\cdot\dot{\gamma},\, \gamma\big)\cdot\frac{1}{\dot{\gamma}}\, \mathrm{d}\gamma,$$

where $\gamma$ is a non-constant differentiable function. Hamilton's Equations thus imply that the Lagrangian does not explicitly depend on the parameter and the Hamiltonian may be set to zero (Beltrami identity). Geometric actions are hence abbreviated with no explicit upper endpoint dependence:

$$\mathcal{A}[q_i] = \int_{t_1}^{t_2} \sum_{i=1}^{n} \frac{\partial L}{\partial \dot{q}_i}\cdot \dot{q}_i\, \mathrm{d}\tau, \qquad \frac{\mathrm{d}S}{\mathrm{d}q_i} = \frac{\partial S}{\partial q_i}.$$

#### 1.4 Lagrangians and Metrics

Henceforth we shall work with the 'real' vectors $\boldsymbol{r}(\tau)$ and omit the generalized (indexed) co-ordinates from the discussion. Segmenting images with variational methods requires carefully defined Lagrangians and actions. Different problems require different Lagrangians, but in two dimensions, they are all of the form

$$L(\boldsymbol{r}, \dot{\boldsymbol{r}}) = \sqrt{\dot{\boldsymbol{r}}\cdot \mathrm{G}\dot{\boldsymbol{r}}}, \tag{3}$$

where G is the metric tensor and $\mathrm{G}\dot{\boldsymbol{r}}$ is a matrix-vector product. Let the metric tensor be isotropic, inhomogeneous and proportional to the identity tensor I. Then the metric tensor, the Lagrangian, and the action become

$$\mathrm{G} = \phi^2(\boldsymbol{r}) \cdot \mathrm{I}, \qquad L = \phi(\boldsymbol{r}) \cdot \|\dot{\boldsymbol{r}}\|, \qquad \mathcal{A}[\boldsymbol{r}] = \int_{t_1}^{t_2} \phi(\boldsymbol{r}) \cdot \|\dot{\boldsymbol{r}}\| \, \mathrm{d}\tau, \tag{4}$$

where $\phi$ is a positive-definite function. Equations (4) imply a geometric action, hence its total rate of change is, owing to the Hamilton-Jacobi Equations:

$$\frac{\mathrm{d}S}{\mathrm{d}\boldsymbol{r}} = \frac{\partial S}{\partial \boldsymbol{r}} = \frac{\partial L}{\partial \dot{\boldsymbol{r}}} = \phi(\boldsymbol{r}) \cdot \frac{\partial \|\dot{\boldsymbol{r}}\|}{\partial \dot{\boldsymbol{r}}} = \phi(\boldsymbol{r}) \cdot \frac{\dot{\boldsymbol{r}}}{\|\dot{\boldsymbol{r}}\|}.$$

Taking the Euclidean norms of both sides, we arrive at the Eikonal Equation

$$\|\boldsymbol{\nabla} S(\boldsymbol{r})\| = \phi(\boldsymbol{r}).$$

### 2 Object Extraction using Minimal Paths

The theoretical framework for the isotropic minimal surface distance map plays the role of the action map and is given by

$$S = \int_a^u \oint_v \Psi(\boldsymbol{R}) \cdot \left\| \frac{\partial \boldsymbol{R}}{\partial v} \right\| \cdot \left\| \frac{\partial \boldsymbol{R}}{\partial u} \right\| \mathrm{d}v \, \mathrm{d}u, \tag{5}$$

as derived in Section 3. A similar distance map is constructed when one considers two closed curves $\boldsymbol{c}_1$ and $\boldsymbol{c}_2$. The path network between them is the set of all contours starting from $\boldsymbol{c}_1$ and ending at $\boldsymbol{c}_2$:

$$N_{\boldsymbol{c}_1}^{\boldsymbol{c}_2} = \left\{ \boldsymbol{R}_{\boldsymbol{c}_1}^q \right\}_{q \in \boldsymbol{c}_2}.$$

An individual path's length is dictated by the enforced metric:

$$D_{\boldsymbol{c}_1}^q = \int_{u(\boldsymbol{c}_1)}^{u(q)} \phi\left(\boldsymbol{R}_{\boldsymbol{c}_1}^q\right) \cdot \left\| \frac{\mathrm{d}\boldsymbol{R}_{\boldsymbol{c}_1}^q}{\mathrm{d}u} \right\| \mathrm{d}u.$$

Upon integrating, we obtain the network's 'energy'

$$S_{\mathrm{net}} = \oint_{q \in \boldsymbol{c}_2} D_{\boldsymbol{c}_1}^q \, \mathrm{d}v = \oint_{q \in \boldsymbol{c}_2} \int_{u(\boldsymbol{c}_1)}^{u(q)} \phi\left(\boldsymbol{R}_{\boldsymbol{c}_1}^q\right) \cdot \left\| \frac{\mathrm{d}\boldsymbol{R}_{\boldsymbol{c}_1}^q}{\mathrm{d}u} \right\| \mathrm{d}u \, \mathrm{d}v \tag{6}$$

which shows striking resemblance with Formula (5). The real difference is the presence of the multiplicative factor $\| \partial \boldsymbol{R} / \partial v \|$. This factor is responsible for preventing the $u =$ const. lines to diverge uncontrollably, since $\| \partial \boldsymbol{R} / \partial v \| \, \mathrm{d}v$ is the infinitesimal distance between such lines.

Despite the similarity of Equations (5) and (6), the theoretical Minimal Surface Problem is fundamentally different because the space of potential solutions is infinite dimensional, as discussed in Section 3. The search for the solution in this infinite dimensional space is not attempted in this paper, rather the *divergence constraint* is implemented which, 'in some degree', imitates the impact of the factor missing from (6).

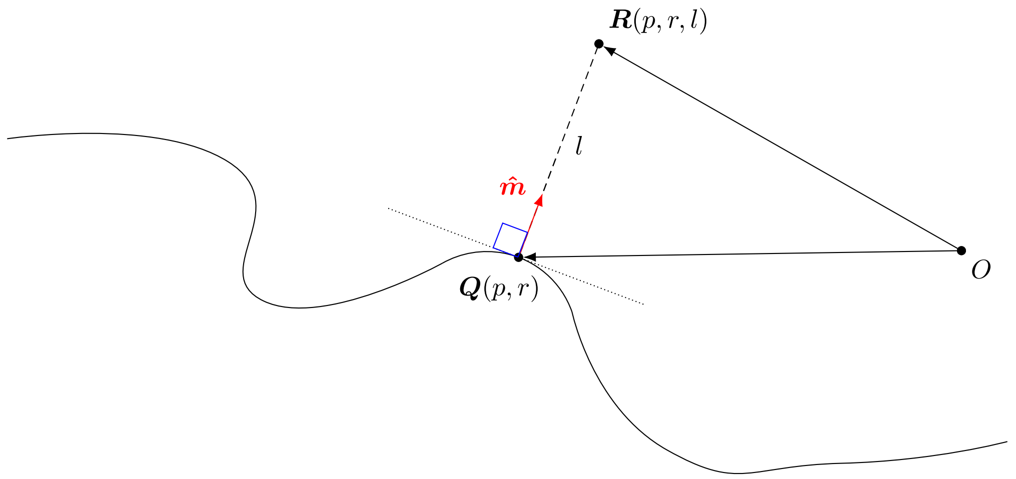


Figure 1: Illustrative sketch of the reference surface co-ordinate system. Points $\boldsymbol{R}(p,r,l)$ are reached from the origin $O$ via the reference surface point $\boldsymbol{Q}(p,r)$ and then the surface unit normal $\hat{\boldsymbol{m}}$. The parameter $l$ is the distance from the surface to the described point.

##### 2.1 The Reference Surface Co-ordinate System

In this section, we discuss the details of the *reference surface co-ordinate system* using the theory of curvilinear co-ordinates. Points $\boldsymbol{R}$ in the 3-dimensional space require 3 co-ordinates $(p,r,l)$ to be uniquely specified. The first 2 of these co-ordinates are sufficient for the description of points $\boldsymbol{Q}$ on a surface which is 2-dimensional. Hence the equation

$$\boldsymbol{R}\big(p,r,l\big) = \boldsymbol{Q}(p,r) + l \cdot \hat{\boldsymbol{m}}(p,r) \tag{7}$$

describes points in 3-dimensional space, where $l$ is their Euclidean distance from the reference surface $\boldsymbol{Q}$ whose unit normal is $\hat{\boldsymbol{m}}$. Hereinafter, vectors with a hat are considered unit length vectors.

Denoting the vector cross product by $\times$, this co-ordinate system is equipped with the basis vectors

$$\boldsymbol{R}_p = \frac{\partial \boldsymbol{R}}{\partial p}, \qquad \boldsymbol{R}_r = \frac{\partial \boldsymbol{R}}{\partial r}, \qquad \boldsymbol{M} = \frac{\partial \boldsymbol{R}}{\partial p} \times \frac{\partial \boldsymbol{R}}{\partial r},$$

and dual basis vectors

$$\boldsymbol{R}^p = \frac{\boldsymbol{R}_r \times \boldsymbol{M}}{\boldsymbol{R}_p \cdot (\boldsymbol{R}_r \times \boldsymbol{M})}, \qquad \boldsymbol{R}^r = \frac{\boldsymbol{M} \times \boldsymbol{R}_p}{\boldsymbol{R}_r \cdot (\boldsymbol{M} \times \boldsymbol{R}_p)}, \qquad \frac{\boldsymbol{M}}{\|\boldsymbol{M}\|^2} = \frac{\boldsymbol{R}_p \times \boldsymbol{R}_r}{\boldsymbol{M} \cdot (\boldsymbol{R}_p \times \boldsymbol{R}_r)}.$$

When $l = 0$, the basis vectors become $\boldsymbol{Q}_p$, $\boldsymbol{Q}_r$, $\boldsymbol{m}$ and the dual basis vectors become $\boldsymbol{Q}^p$, $\boldsymbol{Q}^r$, $\boldsymbol{m}/\|\boldsymbol{m}\|^2$.

**Lemma 2.** *The reference surface co-ordinate system satisfies the following properties.*

1. *Co-linearity:*

$$\boldsymbol{m}(p,r) \times \boldsymbol{M}(p,r,l) = \boldsymbol{0}, \qquad \boldsymbol{m}(p,r) \cdot \boldsymbol{M}(p,r,l) > 0.$$

2. *Divergence:*

$$\hat{\boldsymbol{m}} \cdot \left( \boldsymbol{Q}_p \times \frac{\partial \hat{\boldsymbol{m}}}{\partial r} + \frac{\partial \hat{\boldsymbol{m}}}{\partial p} \times \boldsymbol{Q}_r \right) = \|\boldsymbol{m}\| \cdot (\boldsymbol{\nabla} \cdot \hat{\boldsymbol{m}}).$$

*3. Curvatures:*

$$\boldsymbol{\nabla} \cdot \hat{\boldsymbol{m}} = -K_S, \qquad \hat{\boldsymbol{m}} \cdot \left( \frac{\partial \hat{\boldsymbol{m}}}{\partial p} \times \frac{\partial \hat{\boldsymbol{m}}}{\partial r} \right) = \|\boldsymbol{m}\| \cdot K_G,$$

*where $\boldsymbol{\nabla}\cdot$ is the divergence operator, $K_S$ and $K_G$ are the sum and Gaussian curvatures, respectively.*

*Proof.* Co-linearity arises from the definition of the co-ordinate system (7) and the properties of the scalar and vector triple products. The divergence property follows from the divergence operator expressed in the dual basis [3, pp. 67-68]. The curvature properties follow from [3, pp. 86-91]. □

So far we have only discussed how individual points are described in this new co-ordinate system. Now we shall turn our attention to describing surfaces and their evolution via their normal vectors. Equidistant surfaces from $\boldsymbol{Q}$ are described by the equation

$$\boldsymbol{R}(p, r, l) = \boldsymbol{R}(p, r, \text{const.})$$

and hence their normal vectors $\boldsymbol{M}(p, r, l)$ are co-linear with the reference surface normals $\boldsymbol{m}(p, r)$, satisfying the assertions of Lemma 2. Applying the identities of Lemma 2 and Equation (7) for equidistant surfaces, the magnitude of their normal vector is

$$\begin{aligned}
\|\boldsymbol{M}\| &= \hat{\boldsymbol{m}} \cdot \boldsymbol{M} \\
&= \hat{\boldsymbol{m}} \cdot \left[ \frac{\partial \boldsymbol{R}}{\partial p} \times \frac{\partial \boldsymbol{R}}{\partial r} \right] \\
&= \hat{\boldsymbol{m}} \cdot \left[ \left( \frac{\partial \boldsymbol{Q}}{\partial p} \times \frac{\partial \boldsymbol{Q}}{\partial r} \right) + l \cdot \left( \frac{\partial \boldsymbol{Q}}{\partial p} \times \frac{\partial \hat{\boldsymbol{m}}}{\partial r} + \frac{\partial \hat{\boldsymbol{m}}}{\partial p} \times \frac{\partial \boldsymbol{Q}}{\partial r} \right) + l^2 \cdot \left( \frac{\partial \hat{\boldsymbol{m}}}{\partial p} \times \frac{\partial \hat{\boldsymbol{m}}}{\partial r} \right) \right] \\
&= \hat{\boldsymbol{m}} \cdot \left( \frac{\partial \boldsymbol{Q}}{\partial p} \times \frac{\partial \boldsymbol{Q}}{\partial r} \right) + l \cdot \hat{\boldsymbol{m}} \cdot \left( \frac{\partial \boldsymbol{Q}}{\partial p} \times \frac{\partial \hat{\boldsymbol{m}}}{\partial r} + \frac{\partial \hat{\boldsymbol{m}}}{\partial p} \times \frac{\partial \boldsymbol{Q}}{\partial r} \right) + l^2 \cdot \hat{\boldsymbol{m}} \cdot \left( \frac{\partial \hat{\boldsymbol{m}}}{\partial p} \times \frac{\partial \hat{\boldsymbol{m}}}{\partial r} \right) \\
&= \hat{\boldsymbol{m}} \cdot \boldsymbol{m} + l \cdot \|\boldsymbol{m}\| \cdot \left( \boldsymbol{Q}^r \cdot \frac{\partial \hat{\boldsymbol{m}}}{\partial r} + \boldsymbol{Q}^p \cdot \frac{\partial \hat{\boldsymbol{m}}}{\partial p} \right) + l^2 \cdot \hat{\boldsymbol{m}} \cdot \left( \frac{\partial \hat{\boldsymbol{m}}}{\partial p} \times \frac{\partial \hat{\boldsymbol{m}}}{\partial r} \right) \\
&= \|\boldsymbol{m}\| + l \|\boldsymbol{m}\| \cdot (\boldsymbol{\nabla} \cdot \hat{\boldsymbol{m}}) + l^2 \|\boldsymbol{m}\| \cdot K_{\mathrm{G}} \\
&= \|\boldsymbol{m}\| + l \|\boldsymbol{m}\| \cdot (-K_{\mathrm{S}}) + l^2 \|\boldsymbol{m}\| \cdot K_{\mathrm{G}} \\
&= \|\boldsymbol{m}\| \cdot \left(1 - K_{\mathrm{S}} l + K_{\mathrm{G}} l^2\right).
\end{aligned}$$

Should the distance $l$ from the reference surface become the infinitesimal (and thus equidistant) $\mathrm{d}l$ for the purposes of an update rule, the evolution of the normal vector magnitude reduces to

$$\|\boldsymbol{M}(p, r, \mathrm{d}l)\| = \|\boldsymbol{m}(p, r)\| \cdot (1 - K_{\mathrm{S}} \, \mathrm{d}l), \tag{8}$$

since the second order term becomes negligible. From the area interpretation of the cross product, the evolution of the normal vector is coupled with the evolution of the surface patch area.

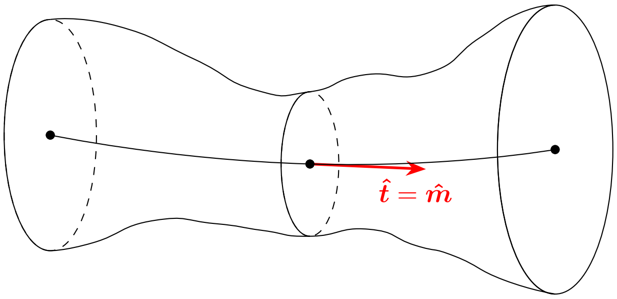


Figure 2: Surface evolution along an extremal path. The coinciding unit tangent and unit normal is the unique property of the isotropic metric.

#### 2.2 Evolution of an Area Element

Assuming constant $(p, r)$, the magnitude of the surface normal is only a function of $l$. By definition, $\boldsymbol{m}(p, r) = \boldsymbol{M}(p, r, 0)$ and the update rule (8) becomes, after rearranging and suppressing the $(p, r)$ dependence,

$$\frac{\|\boldsymbol{M}(\mathrm{d}l)\|}{\|\boldsymbol{M}(0)\|} = 1 - K_{\mathrm{S}}\,\mathrm{d}l \qquad \Rightarrow \qquad \frac{\|\boldsymbol{M}(S + \mathrm{d}S)\|}{\|\boldsymbol{M}(S)\|} = 1 - K_{\mathrm{S}}\frac{\mathrm{d}l}{\mathrm{d}S}\,\mathrm{d}S,$$

using the chain rule and exploiting the fact that $l = 0$ is an arbitrarily chosen starting point of the evolution. Assuming that the derivative $\mathrm{d}l/\mathrm{d}S$ is known, the evolution equation with respect to the action values is complete.

The length of one-dimensional elements inside an elementary surface patch is, 'in some average sense', proportional to the square rooted area of that patch. We are going to see this in more detail in Subsection 2.3 where this approximate proportionality will correspond to leaving out higher order terms from the Taylor expansion of the action. The exact rate of change of vector magnitudes is the function of the direction tangential to the plane in which the one-dimensional element passes. 'In some average sense' is our best description as far as local length information is concerned, should we wish to keep the isotropic nature of our equations. Using the binomial approximation for small sum curvatures $K_{\mathrm{S}}$, the rate of change of the length of the 'mean elementary distances' is

$$\sqrt{\frac{\|\boldsymbol{M}(S + \mathrm{d}S)\|}{\|\boldsymbol{M}(S)\|}} = \sqrt{1 - K_{\mathrm{S}}\frac{\mathrm{d}l}{\mathrm{d}S}} \approx 1 - \frac{1}{2}K_{\mathrm{S}}\frac{\mathrm{d}l}{\mathrm{d}S}. \tag{9}$$

#### 2.3 The Divergence Constraint

The evolution equation

$$\frac{\mathrm{d}S}{\mathrm{d}\boldsymbol{r}} = \phi(\boldsymbol{r}) \cdot \frac{\dot{\boldsymbol{r}}}{\|\dot{\boldsymbol{r}}\|} \tag{10}$$

describes how the distance map is built from an initial $S = 0$ surface in the $\dot{\boldsymbol{r}}/\|\dot{\boldsymbol{r}}\|$ direction. Once a set $S =$ const. is known, the updated constant value is $S + \mathrm{d}S$ based on the evolution equation. Since the tangents of the minimal paths coincide with the unit normals $\hat{\boldsymbol{m}}$ of the minimal surfaces in the isotropic case, the update rule can be expressed only with respect to the normal distance. Project the evolution equation (10) onto the unit normal vector:

$$\frac{\mathrm{d}S}{\mathrm{d}l} = \frac{\mathrm{d}S}{\mathrm{d}\boldsymbol{r}} \boldsymbol{\cdot} \hat{\boldsymbol{m}} = \phi(\boldsymbol{r}) \cdot \frac{\dot{\boldsymbol{r}}}{\|\dot{\boldsymbol{r}}\|} \boldsymbol{\cdot} \hat{\boldsymbol{m}} = \phi(\boldsymbol{r}) \cdot \hat{\boldsymbol{m}} \boldsymbol{\cdot} \hat{\boldsymbol{m}} = \phi(\boldsymbol{r}),$$

from which the normal infinitesimal elongation is found to first order:

$$\mathrm{d}l = \frac{\mathrm{d}S}{\phi(\boldsymbol{r})}. \tag{11}$$

In order to incorporate the curvature term into our updating scheme, we consider the Taylor expansion of the Hamilton's principal function up to second order, involving the Laplacian operator and the big $\mathcal{O}$ notation:

$$\begin{aligned}
S(\boldsymbol{r}+\mathrm{d}\boldsymbol{r}) - S(\boldsymbol{r}) &= \mathrm{d}\boldsymbol{r} \boldsymbol{\cdot} \frac{\mathrm{d}S}{\mathrm{d}\boldsymbol{r}} + \frac{1}{2}\|\mathrm{d}\boldsymbol{r}\|^2 \cdot \frac{\mathrm{d}^2 S}{\mathrm{d}\boldsymbol{r}^2} + \mathcal{O}\Big(\|\mathrm{d}\boldsymbol{r}\|^3\Big) \\
&= \mathrm{d}\boldsymbol{r} \boldsymbol{\cdot} (\phi(\boldsymbol{r}) \cdot \hat{\boldsymbol{m}}) + \frac{1}{2}\|\mathrm{d}\boldsymbol{r}\|^2 \cdot \boldsymbol{\nabla} \boldsymbol{\cdot} \boldsymbol{\nabla} S(\boldsymbol{r}) + \mathcal{O}\Big(\|\mathrm{d}\boldsymbol{r}\|^3\Big) \\
&= \phi(\boldsymbol{r}) \cdot (\mathrm{d}\boldsymbol{r} \boldsymbol{\cdot} \hat{\boldsymbol{m}}) + \frac{1}{2}\|\mathrm{d}\boldsymbol{r}\|^2 \cdot \boldsymbol{\nabla} \boldsymbol{\cdot} (\phi \cdot \hat{\boldsymbol{m}})(\boldsymbol{r}) + \mathcal{O}\Big(\|\mathrm{d}\boldsymbol{r}\|^3\Big) \\
&= \phi(\boldsymbol{r}) \cdot (\mathrm{d}\boldsymbol{r} \boldsymbol{\cdot} \hat{\boldsymbol{m}}) + \frac{1}{2}\|\mathrm{d}\boldsymbol{r}\|^2 \cdot (\boldsymbol{\nabla}\phi(\boldsymbol{r}) \boldsymbol{\cdot} \hat{\boldsymbol{m}} + \phi(\boldsymbol{r}) \cdot \boldsymbol{\nabla} \boldsymbol{\cdot} \hat{\boldsymbol{m}}) + \mathcal{O}\Big(\|\mathrm{d}\boldsymbol{r}\|^3\Big) \\
&= \phi(\boldsymbol{r}) \cdot (\mathrm{d}\boldsymbol{r} \boldsymbol{\cdot} \hat{\boldsymbol{m}}) + \frac{1}{2}\|\mathrm{d}\boldsymbol{r}\|^2 \cdot \left(\boldsymbol{\nabla}\phi(\boldsymbol{r}) \boldsymbol{\cdot} \frac{\dot{\boldsymbol{r}}}{\|\dot{\boldsymbol{r}}\|} - \phi(\boldsymbol{r}) \cdot K_{\mathrm{S}}\right) + \mathcal{O}\Big(\|\mathrm{d}\boldsymbol{r}\|^3\Big) \\
&= \phi(\boldsymbol{r}) \cdot (\mathrm{d}\boldsymbol{r} \boldsymbol{\cdot} \hat{\boldsymbol{m}}) + \frac{1}{2}\|\mathrm{d}\boldsymbol{r}\|^2 \cdot (0 - \phi(\boldsymbol{r}) \cdot K_{\mathrm{S}}) + \mathcal{O}\Big(\|\mathrm{d}\boldsymbol{r}\|^3\Big) \\
&= \phi(\boldsymbol{r}) \cdot (\mathrm{d}\boldsymbol{r} \boldsymbol{\cdot} \hat{\boldsymbol{m}}) - \frac{1}{2}\|\mathrm{d}\boldsymbol{r}\|^2 \phi(\boldsymbol{r}) \cdot K_{\mathrm{S}} + \mathcal{O}\Big(\|\mathrm{d}\boldsymbol{r}\|^3\Big) \\
&= \phi(\boldsymbol{r}) \cdot \left((\mathrm{d}\boldsymbol{r} \boldsymbol{\cdot} \hat{\boldsymbol{m}}) - \frac{1}{2}\|\mathrm{d}\boldsymbol{r}\|^2 K_{\mathrm{S}}\right) + \mathcal{O}\Big(\|\mathrm{d}\boldsymbol{r}\|^3\Big)
\end{aligned}$$

because when expanding the divergence, only the curvature term contributes, as the normal and tangential vectors of the scalar field $\phi$ are perpendicular to each other. Let us now consider infinitesimal advances in the direction of the normal vector, i.e. $\mathrm{d}\boldsymbol{r} = \mathrm{d}l \cdot \hat{\boldsymbol{m}}$ and ignore the third and higher order contributions. Then the Taylor expansion reduces to

$$\mathrm{d}S = \phi(\boldsymbol{r})\,\mathrm{d}l \cdot \left(1 - \frac{1}{2} K_{\mathrm{S}}\,\mathrm{d}l\right), \tag{12}$$

in accordance with the rate of change of 'mean elementary distances' formula (9). Equation (12) is a quadratic in $\mathrm{d}l$ and solved by

$$\mathrm{d}l = \frac{1 - \sqrt{1 - 4K_{\mathrm{S}} \cdot \mathrm{d}S/\phi(\boldsymbol{r})}}{2K_{\mathrm{S}}}. \tag{13}$$

The sign chosen for the square root is guided by the principle that in the limit of zero sum curvature, the infinitesimal normal dilation reduces to the first order formula (11). Using L'Hôpital's rule, write

$$\lim_{K_{\mathrm{S}}\to 0} \mathrm{d}l = \lim_{K_{\mathrm{S}}\to 0} \frac{1 - \sqrt{1 - 4K_{\mathrm{S}} \cdot \mathrm{d}S/\phi(\boldsymbol{r})}}{2K_{\mathrm{S}}} = \lim_{K_{\mathrm{S}}\to 0} -\frac{1}{2} \cdot \frac{-4 \cdot \mathrm{d}S/\phi(\boldsymbol{r})}{\sqrt{1 - 4K_{\mathrm{S}} \cdot \mathrm{d}S/\phi(\boldsymbol{r})}} \cdot \frac{1}{2} = \frac{\mathrm{d}S}{\phi(\boldsymbol{r})},$$

as expected. We must, of course, account for the possibility that the discriminant of the quadratic (12) becomes negative. In order to prevent this, we demand

$$\max \mathrm{d}S = \frac{\min \phi}{4K_{\mathrm{S}}} \qquad \Rightarrow \qquad \max \mathrm{d}l = \frac{1}{2K_{\mathrm{S}}}.$$

Another method of applying the second order update rule (13) is to let $\mathrm{d}S = \text{const.}$ and use the minimum value of the scalar field $\phi$ allowed by the first order update rule (11). In this case, the maximum allowed normal dilation is

$$\max \mathrm{d}l = \frac{1}{2K_\mathrm{S}} = \frac{1}{2 \cdot \min \phi / (4 \cdot \max \mathrm{d}S)} = \frac{1}{2 \cdot \min \phi / (4 \cdot \mathrm{d}S)} = \frac{2\,\mathrm{d}S}{\min \phi}.$$

The enforced metric is

$$\phi(\boldsymbol{r}) = \alpha + (1 - \alpha) \cdot \exp(-\beta \|\boldsymbol{\nabla} I(\boldsymbol{r})\|),$$

where $\alpha$ and $\beta$ are parameters set by the user. Here, $I$ stands for the image intensity function restricted to the $[0, 1]$ interval.

### 3 Theoretical Minimal Surface Framework

In the 3-dimensional Euclidean space, points $\boldsymbol{R}$ on surfaces are parameterized by functions of two co-ordinates: $\boldsymbol{R} = \boldsymbol{R}(u, v)$. The surface $\Sigma$ between the two contours is a set of points such that the parameter values are bounded:

$$\Sigma = \{\boldsymbol{R}(u, v) : \; u_1 \le u \le u_2 \quad \text{and} \quad v_1 \le v \le v_2\}.$$

We may choose a parameterization such that all the bounds are constant in an appropriate co-ordinate system. Consider the first order perturbation of the surface points in the direction of a non-zero arbitrary vector $\boldsymbol{h}$:

$$\boldsymbol{R} \to \boldsymbol{R} + \varepsilon \boldsymbol{h},$$

where $\varepsilon$ is the perturbation parameter. The surface normal is

$$\boldsymbol{N} = \frac{\partial \boldsymbol{R}}{\partial u} \times \frac{\partial \boldsymbol{R}}{\partial v},$$

and its first order perturbation satisfies

$$\begin{aligned}
\boldsymbol{N} + \delta \boldsymbol{N} + \mathcal{O}(\varepsilon^2) &= \frac{\partial(\boldsymbol{R} + \varepsilon \boldsymbol{h})}{\partial u} \times \frac{\partial(\boldsymbol{R} + \varepsilon \boldsymbol{h})}{\partial v} + \mathcal{O}(\varepsilon^2) \\
&= \left(\frac{\partial \boldsymbol{R}}{\partial u} + \varepsilon \frac{\partial \boldsymbol{h}}{\partial u}\right) \times \left(\frac{\partial \boldsymbol{R}}{\partial v} + \varepsilon \frac{\partial \boldsymbol{h}}{\partial v}\right) + \mathcal{O}(\varepsilon^2) \\
&= \frac{\partial \boldsymbol{R}}{\partial u} \times \frac{\partial \boldsymbol{R}}{\partial v} + \varepsilon \left(\frac{\partial \boldsymbol{R}}{\partial u} \times \frac{\partial \boldsymbol{h}}{\partial v} + \frac{\partial \boldsymbol{h}}{\partial u} \times \frac{\partial \boldsymbol{R}}{\partial v}\right) + \mathcal{O}(\varepsilon^2) \\
&= \boldsymbol{N} + \varepsilon \left(\frac{\partial \boldsymbol{R}}{\partial u} \times \frac{\partial \boldsymbol{h}}{\partial v} + \frac{\partial \boldsymbol{h}}{\partial u} \times \frac{\partial \boldsymbol{R}}{\partial v}\right) + \mathcal{O}(\varepsilon^2),
\end{aligned}$$

revealing the first variation in the normal vector

$$\delta \boldsymbol{N} = \varepsilon \left(\frac{\partial \boldsymbol{R}}{\partial u} \times \frac{\partial \boldsymbol{h}}{\partial v} + \frac{\partial \boldsymbol{h}}{\partial u} \times \frac{\partial \boldsymbol{R}}{\partial v}\right).$$

Assuming geometric functionals, the surface action is

$$\mathcal{A}[\boldsymbol{R}] = \iint_\Sigma L(\boldsymbol{R}, \boldsymbol{N}) \,\mathrm{d}u \,\mathrm{d}v.$$

While its functional differential, expanding to first order in $\varepsilon$,

$$
\begin{aligned}
\delta\mathcal{A} &= \iint_\Sigma L\left(\boldsymbol{R}+\varepsilon\boldsymbol{h}, \boldsymbol{N}+\delta\boldsymbol{N}\right) - L\left(\boldsymbol{R},\boldsymbol{N}\right)\mathrm{d}u\,\mathrm{d}v \\
&= \iint_\Sigma \frac{\partial L}{\partial \boldsymbol{R}}\cdot\varepsilon\boldsymbol{h} + \frac{\partial L}{\partial \boldsymbol{N}}\cdot\delta\boldsymbol{N}\,\mathrm{d}u\,\mathrm{d}v \\
&= \iint_\Sigma \frac{\partial L}{\partial \boldsymbol{R}}\cdot\varepsilon\boldsymbol{h} + \frac{\partial L}{\partial \boldsymbol{N}}\cdot\varepsilon\left(\frac{\partial\boldsymbol{R}}{\partial u}\times\frac{\partial\boldsymbol{h}}{\partial v}+\frac{\partial\boldsymbol{h}}{\partial u}\times\frac{\partial\boldsymbol{R}}{\partial v}\right)\mathrm{d}u\,\mathrm{d}v \\
&= \iint_\Sigma \varepsilon\boldsymbol{h}\cdot\frac{\partial L}{\partial \boldsymbol{R}} + \varepsilon\frac{\partial\boldsymbol{h}}{\partial u}\cdot\left(\frac{\partial\boldsymbol{R}}{\partial v}\times\frac{\partial L}{\partial\boldsymbol{N}}\right) + \varepsilon\frac{\partial\boldsymbol{h}}{\partial v}\cdot\left(\frac{\partial L}{\partial\boldsymbol{N}}\times\frac{\partial\boldsymbol{R}}{\partial u}\right)\mathrm{d}u\,\mathrm{d}v \\
&= \iint_\Sigma \varepsilon\boldsymbol{h}\cdot\frac{\partial L}{\partial \boldsymbol{R}}\,\mathrm{d}u\,\mathrm{d}v + \iint_\Sigma \varepsilon\frac{\partial\boldsymbol{h}}{\partial u}\cdot\left(\frac{\partial\boldsymbol{R}}{\partial v}\times\frac{\partial L}{\partial\boldsymbol{N}}\right)\mathrm{d}u\,\mathrm{d}v + \iint_\Sigma \varepsilon\frac{\partial\boldsymbol{h}}{\partial v}\cdot\left(\frac{\partial L}{\partial\boldsymbol{N}}\times\frac{\partial\boldsymbol{R}}{\partial u}\right)\mathrm{d}u\,\mathrm{d}v, \qquad (14)
\end{aligned}
$$

using the cyclic property of the scalar triple product. We now perform integration by parts on the second and third terms to move the derivatives away from $\boldsymbol{h}$. Write

$$
\iint_\Sigma \varepsilon\frac{\partial\boldsymbol{h}}{\partial u}\cdot\left(\frac{\partial\boldsymbol{R}}{\partial v}\times\frac{\partial L}{\partial\boldsymbol{N}}\right)\mathrm{d}u\,\mathrm{d}v = \int_{v_1}^{v_2}\left[\varepsilon\boldsymbol{h}\cdot\left(\frac{\partial\boldsymbol{R}}{\partial v}\times\frac{\partial L}{\partial\boldsymbol{N}}\right)\right]_{u_1}^{u_2}\mathrm{d}v - \iint_\Sigma \varepsilon\boldsymbol{h}\cdot\frac{\partial}{\partial u}\left(\frac{\partial\boldsymbol{R}}{\partial v}\times\frac{\partial L}{\partial\boldsymbol{N}}\right)\mathrm{d}u\,\mathrm{d}v,
$$

$$
\iint_\Sigma \varepsilon\frac{\partial\boldsymbol{h}}{\partial v}\cdot\left(\frac{\partial L}{\partial\boldsymbol{N}}\times\frac{\partial\boldsymbol{R}}{\partial u}\right)\mathrm{d}u\,\mathrm{d}v = \int_{u_1}^{u_2}\left[\varepsilon\boldsymbol{h}\cdot\left(\frac{\partial L}{\partial\boldsymbol{N}}\times\frac{\partial\boldsymbol{R}}{\partial u}\right)\right]_{v_1}^{v_2}\mathrm{d}u - \iint_\Sigma \varepsilon\boldsymbol{h}\cdot\frac{\partial}{\partial v}\left(\frac{\partial L}{\partial\boldsymbol{N}}\times\frac{\partial\boldsymbol{R}}{\partial u}\right)\mathrm{d}u\,\mathrm{d}v.
$$

The double integrals are then absorbed into one double integral, assumed to vanish at the extremal surface:

$$
\iint_\Sigma \varepsilon\boldsymbol{h}\cdot\left[\frac{\partial L}{\partial\boldsymbol{R}} - \frac{\partial}{\partial u}\left(\frac{\partial\boldsymbol{R}}{\partial v}\times\frac{\partial L}{\partial\boldsymbol{N}}\right) - \frac{\partial}{\partial v}\left(\frac{\partial L}{\partial\boldsymbol{N}}\times\frac{\partial\boldsymbol{R}}{\partial u}\right)\right]\mathrm{d}u\,\mathrm{d}v = 0.
$$

Since $\varepsilon\boldsymbol{h}$ was a non-zero arbitrary vector, the integral vanishes if and only if the Euler-Lagrange Equation for extremal surfaces is satisfied:

$$
\frac{\partial L}{\partial\boldsymbol{R}} - \frac{\partial}{\partial u}\left(\frac{\partial\boldsymbol{R}}{\partial v}\times\frac{\partial L}{\partial\boldsymbol{N}}\right) - \frac{\partial}{\partial v}\left(\frac{\partial L}{\partial\boldsymbol{N}}\times\frac{\partial\boldsymbol{R}}{\partial u}\right) = \boldsymbol{0}.
$$

Continuing with the assumption that the Euler-Lagrange Equation is satisfied, the variation of the action now only contains the single integrals and it becomes the infinitesimal change in the Hamilton's principal function. Let $\varepsilon\boldsymbol{h} = \delta\boldsymbol{R}$ and write (14), owing to Green's theorem and Stokes' theorem,

$$
\begin{aligned}
\mathrm{d}S &= \int_{v_1}^{v_2}\left[\delta\boldsymbol{R}\cdot\left(\frac{\partial\boldsymbol{R}}{\partial v}\times\frac{\partial L}{\partial\boldsymbol{N}}\right)\right]_{u_1}^{u_2}\mathrm{d}v + \int_{u_1}^{u_2}\left[\delta\boldsymbol{R}\cdot\left(\frac{\partial L}{\partial\boldsymbol{N}}\times\frac{\partial\boldsymbol{R}}{\partial u}\right)\right]_{v_1}^{v_2}\mathrm{d}u \\
&= \iint_\Sigma \frac{\partial}{\partial u}\left[\delta\boldsymbol{R}\cdot\left(\frac{\partial\boldsymbol{R}}{\partial v}\times\frac{\partial L}{\partial\boldsymbol{N}}\right)\right]\mathrm{d}u\,\mathrm{d}v + \iint_\Sigma \frac{\partial}{\partial v}\left[\delta\boldsymbol{R}\cdot\left(\frac{\partial L}{\partial\boldsymbol{N}}\times\frac{\partial\boldsymbol{R}}{\partial u}\right)\right]\mathrm{d}u\,\mathrm{d}v \\
&= \iint_\Sigma \frac{\partial}{\partial u}\left[\delta\boldsymbol{R}\cdot\left(\frac{\partial\boldsymbol{R}}{\partial v}\times\frac{\partial L}{\partial\boldsymbol{N}}\right)\right] + \frac{\partial}{\partial v}\left[\delta\boldsymbol{R}\cdot\left(\frac{\partial L}{\partial\boldsymbol{N}}\times\frac{\partial\boldsymbol{R}}{\partial u}\right)\right]\mathrm{d}u\,\mathrm{d}v \\
&= \iint_\Sigma \frac{\partial}{\partial u}\left[\left(\frac{\partial L}{\partial\boldsymbol{N}}\times\delta\boldsymbol{R}\right)\cdot\frac{\partial\boldsymbol{R}}{\partial v}\right] - \frac{\partial}{\partial v}\left[\left(\frac{\partial L}{\partial\boldsymbol{N}}\times\delta\boldsymbol{R}\right)\cdot\frac{\partial\boldsymbol{R}}{\partial u}\right]\mathrm{d}u\,\mathrm{d}v \\
&= \oint_\gamma \left(\frac{\partial L}{\partial\boldsymbol{N}}\times\delta\boldsymbol{R}\right)\cdot\mathrm{d}\boldsymbol{B}, \qquad (15)
\end{aligned}
$$

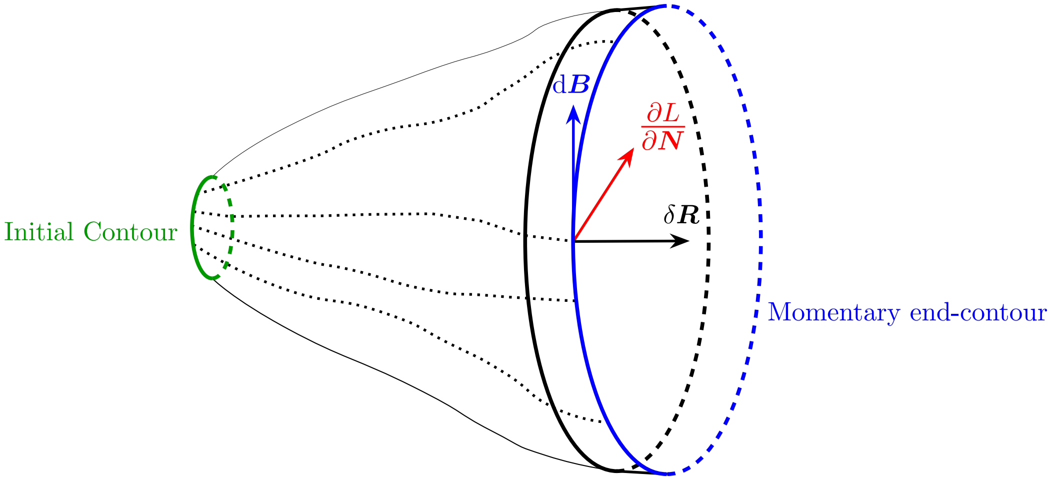


Figure 3: Distance map building based on the parallelepiped spanned by the vectors $\partial L/\partial \boldsymbol{N}$, $\delta\boldsymbol{R}$ and $\mathrm{d}\boldsymbol{B}$. The paralellepiped volumes are summed over the momentary end-contour $\gamma$, yielding the infinitesimal change in the Hamilton's principal function.

where $\boldsymbol{B}$ stands for the points on the varying end boundary closed curve $\gamma$.

Choosing the special parameterization $u = \text{const.}$, the differential formulation of the Eikonal Equation (15) becomes

$$\mathrm{d}S = \oint_v \left(\frac{\partial L}{\partial \boldsymbol{N}} \times \delta\boldsymbol{R}\right) \cdot \frac{\partial \boldsymbol{R}}{\partial v}\,\mathrm{d}v = \oint_v \left(\frac{\partial \boldsymbol{R}}{\partial v} \times \frac{\partial L}{\partial \boldsymbol{N}}\right) \cdot \delta\boldsymbol{R}\,\mathrm{d}v$$

because the points $\boldsymbol{B}$ depend only on $v$ in this case. Analogously to the Hamilton-Jacobi Equations (2), we can nominate the action gradient and the inner product

$$\boldsymbol{\nabla} S = \frac{\partial \boldsymbol{R}}{\partial v} \times \frac{\partial L}{\partial \boldsymbol{N}}, \qquad \langle \boldsymbol{\nabla} S, \delta\boldsymbol{R}\rangle = \oint_v \boldsymbol{\nabla} S \cdot \delta\boldsymbol{R}\,\mathrm{d}v,$$

forming a triplet-space. The Minimal Surface Problem thus reduces to the task of finding minimal paths in the space of boundary contours, see Figure 3. Had we chosen a different parameterization with the same information content, our argument would still be valid. By the definition of the cross product, $\boldsymbol{\nabla} S$ is perpendicular to the momentary contour points. Thus, the parallelity of the normal and the variation reduces the integrand to the product of the norms:

$$\boldsymbol{\nabla} S \parallel \delta\boldsymbol{R} \qquad \Rightarrow \qquad \mathrm{d}S = \oint_v \left\|\frac{\partial \boldsymbol{R}}{\partial v} \times \frac{\partial L}{\partial \boldsymbol{N}}\right\| \cdot \|\delta\boldsymbol{R}\|\,\mathrm{d}v$$

which is a formula suitable for the action map construction.

#### 3.1 Application to the Isotropic Case

Similarly to Equation (3), the Lagrangian is of the form

$$L(\boldsymbol{R}, \boldsymbol{N}) = \sqrt{\boldsymbol{N} \cdot \mathrm{G_T}\boldsymbol{N}},$$

where $\boldsymbol{N}$ is the normal vector of the tangent plane whose metric tensor is $\mathrm{G_T}$. Let the metric tensor be isotropic, inhomogeneous and proportional to the identity tensor I. Then the metric tensor, the Lagrangian, and the action become

$$\mathrm{G_T} = \Psi^2(\boldsymbol{R}) \cdot \mathrm{I}, \qquad L = \Psi(\boldsymbol{R}) \cdot \|\boldsymbol{N}\|, \qquad \mathcal{A}[\boldsymbol{R}] = \iint_\Sigma \Psi(\boldsymbol{R}) \cdot \|\boldsymbol{N}\| \,\mathrm{d}u \,\mathrm{d}v,$$

where $\Psi$ is a positive-definite function. Using the area interpretation of the cross product, the Hamilton's principal function of the problem becomes either

$$S = \iint_\Sigma \Psi(\boldsymbol{R}) \cdot \|\boldsymbol{N}\| \,\mathrm{d}u \,\mathrm{d}v = \iint_\Sigma \Psi(\boldsymbol{R}) \cdot \frac{\boldsymbol{N}}{\|\boldsymbol{N}\|} \boldsymbol{\cdot} \boldsymbol{N} \,\mathrm{d}u \,\mathrm{d}v = \iint_\Sigma \Psi(\boldsymbol{R}) \cdot \frac{\boldsymbol{N}}{\|\boldsymbol{N}\|} \boldsymbol{\cdot} \mathrm{d}\boldsymbol{A},$$

or, operating under the assumption $\boldsymbol{\nabla} S \parallel \delta\boldsymbol{R}$, its differential form,

$$\mathrm{d}S = \oint_v \left\| \frac{\partial \boldsymbol{R}}{\partial v} \times \frac{\partial L}{\partial \boldsymbol{N}} \right\| \cdot \|\delta\boldsymbol{R}\| \,\mathrm{d}v = \oint_v \left\| \frac{\partial \boldsymbol{R}}{\partial v} \times \left( \Psi(\boldsymbol{R}) \cdot \frac{\boldsymbol{N}}{\|\boldsymbol{N}\|} \right) \right\| \cdot \|\delta\boldsymbol{R}\| \,\mathrm{d}v = \oint_v \Psi(\boldsymbol{R}) \cdot \left\| \frac{\partial \boldsymbol{R}}{\partial v} \right\| \cdot \|\delta\boldsymbol{R}\| \,\mathrm{d}v$$

which can be integrated to give

$$S = \int_a^u \oint_v \Psi(\boldsymbol{R}) \cdot \left\| \frac{\partial \boldsymbol{R}}{\partial v} \right\| \cdot \left\| \frac{\partial \boldsymbol{R}}{\partial u} \right\| \mathrm{d}v \,\mathrm{d}u,$$

with an arbitrary lower bound, since it can be absorbed by $S$.

### 4 Minimal Contour Framework

In the Minimal Contour Problem, the theoretical background is the same as discussed in the Lagrangian and Metrics Subsection 1.4. In this case, the enforced metric is

$$\phi(\boldsymbol{r}) = (\boldsymbol{\nabla} I \boldsymbol{\cdot} \hat{\boldsymbol{n}}) \cdot \frac{1 - \exp\left(-\beta \cdot \frac{\|\boldsymbol{\nabla} I\|}{\max(\|\boldsymbol{\nabla} I\|)}\right)}{\|\boldsymbol{\nabla} I\|} + \sqrt{1 - \left(\frac{\partial I}{\partial y}\hat{n}_x - \frac{\partial I}{\partial x}\hat{n}_y\right)^2 \cdot \left(\frac{1 - \exp\left(-\beta \cdot \frac{\|\boldsymbol{\nabla} I\|}{\max(\|\boldsymbol{\nabla} I\|)}\right)}{\|\boldsymbol{\nabla} I\|}\right)^2},$$

where $\boldsymbol{\nabla} I$ is the image intensity gradient and $\hat{\boldsymbol{n}}$ is the unit normal of the momentary contour whose Cartesian components are $\hat{n}_x, \hat{n}_y$.

### 5 Mean contour framework

In this model, we consider simple, planar curves to be closed, continuous, one-parameter ($t \in [0, T]$) objects. From now on, we refer to them as 'contours'. The principal representations of contours are often given by a position vector with respect to a standard basis $\mathbf{i}$, $\mathbf{j}$ as $\boldsymbol{r}(t) = x(t)\mathbf{i} + y(t)\mathbf{j}$, $\boldsymbol{r}(0) = \boldsymbol{r}(T)$, where $x(t)$ and $y(t)$ are the coordinate functions of the contour. To get a parametrization-invariant representation of the contours, one possibility is to get the square root velocity (SRV) representation of them, however, it does not retain relative translation information, so it cannot be used for annotation purposes. Thus, we modify this representation and apply the rescaled position by square root velocity (RPSV) function:

$$\boldsymbol{q}(t) = \boldsymbol{r}(t)\sqrt{\|\dot{\boldsymbol{r}}(t)\|}.$$

Note, that $\boldsymbol{q}(t)$ and $\boldsymbol{r}(t)$ lie in the same direction, so reconstructing the original contour only requires us to get the $\|\boldsymbol{r}(t)\|$ values at every parameter value $t$.

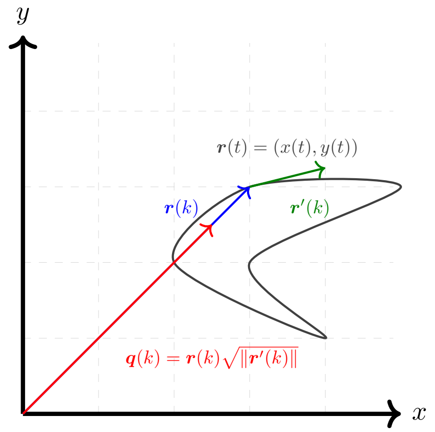


Figure 4: Visualization of the RPSV representation of an arbitrary point $k \in [0, T]$. As can be seen, the direction of $\boldsymbol{q}(k)$ is the same as the direction of $\boldsymbol{r}(k)$.

In this representation space, the distance function is invariant with respect to the reparametrization of the points ($d^2(\boldsymbol{q}_1, \boldsymbol{q}_2) = d^2(\boldsymbol{q}_1 \circ \gamma, \boldsymbol{q}_2 \circ \gamma)$), so we can now calculate the mean of several contours (which, in our case, are annotations), as it will be parametrization-invariant:

$$\boldsymbol{r}(t)\sqrt{\|\dot{\boldsymbol{r}}(t)\|} = \frac{1}{n}\sum_{i=1}^{n} \boldsymbol{r}_i(t_i)\sqrt{\|\dot{\boldsymbol{r}}(t_i)\|},$$

where $n$ is the number of annotations, and $t_i = \gamma_i(t)$ is the parametrization of the $i$-th contour.

To find the optimal mean contour of the contour system, we have to modify the $\gamma_i$ values for the contours to be as close as possible to the mean contour in the representation space. Formally, this means finding a reparameterization function $\gamma_i$ for every contour $i$ that minimizes the objective function

$$\sum_{i=1}^{n} \oint \left(\boldsymbol{r}(t)\sqrt{\|\dot{\boldsymbol{r}}(t)\|} - \boldsymbol{r}_i(t_i)\sqrt{\|\dot{\boldsymbol{r}}_i(t_i)\|}\right)^2 \mathrm{d}t, \tag{16}$$

where $\boldsymbol{r}(t)$ is the mean contour according to the "current" reparameterizations. The solution to this minimalization problem is equivalent to finding the pairwise minimal distance to a reference contour (say $\boldsymbol{r}_1$):

$$\min_{\gamma_k} \oint \left(\boldsymbol{r}_1(t)\sqrt{\|\dot{\boldsymbol{r}}_1(t)\|} - \boldsymbol{r}_k(t_k)\sqrt{\|\dot{\boldsymbol{r}}_k(t_k)\|}\right)^2 \mathrm{d}t.$$

This minimization problem can now be solved with the tools of variational calculus, where the Lagrangian and its derivatives are:

$$\begin{aligned}
L(\gamma_k, \dot{\gamma}_k) &= \left(\boldsymbol{r}_1\sqrt{\|\dot{\boldsymbol{r}}_1\|} - \boldsymbol{r}_k\sqrt{\|\boldsymbol{r}'_k\|\dot{\gamma}_k}\right)^2 \\
\frac{\partial L}{\partial \gamma_k} &= -2\left(\boldsymbol{r}_1\sqrt{\|\dot{\boldsymbol{r}}_1\|} - \boldsymbol{r}_k\sqrt{\|\boldsymbol{r}'_k\|\dot{\gamma}_k}\right) \cdot \left(\boldsymbol{r}'_k\sqrt{\|\boldsymbol{r}'_k\|\dot{\gamma}_k} + \boldsymbol{r}_k\frac{\dot{\gamma}_k \boldsymbol{e}_k \cdot \boldsymbol{r}''_k}{2\sqrt{\|\boldsymbol{r}'_k\|\dot{\gamma}_k}}\right) \\
\frac{\partial L}{\partial \dot{\gamma}_k} &= -\left(\boldsymbol{r}_1\sqrt{\|\dot{\boldsymbol{r}}_1\|} - \boldsymbol{r}_k\sqrt{\|\boldsymbol{r}'_k\|\dot{\gamma}_k}\right) \cdot \boldsymbol{r}_k\frac{\|\boldsymbol{r}'_k\|}{\sqrt{\|\boldsymbol{r}'_k\|\dot{\gamma}_k}}.
\end{aligned} \tag{17}$$

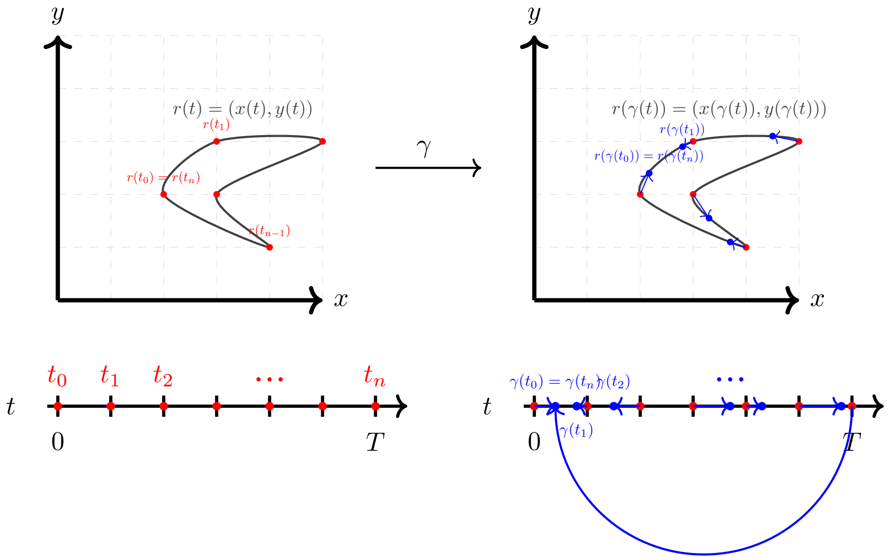


Figure 5: The arc-length parametrization of the contour $\boldsymbol{r}(t)$ (left), and the effect of a reparametrization $\gamma$ (right).

From the relations between the differential operators

$$\begin{aligned}
\frac{\mathrm{d}}{\mathrm{d}\gamma} &= \frac{1}{\dot\gamma}\frac{\mathrm{d}}{\mathrm{d}t} \\
\frac{\mathrm{d}^2}{\mathrm{d}\gamma^2} &= \frac{1}{\dot\gamma}\left(-\frac{\ddot\gamma}{\dot\gamma^2}\frac{\mathrm{d}}{\mathrm{d}t} + \frac{1}{\dot\gamma}\frac{\mathrm{d}^2}{\mathrm{d}t^2}\right),
\end{aligned} \tag{18}$$

the Euler-Lagrange equation for the $k$-th diffeomorphism $\gamma_k = \gamma_k\,(t)$ is:

$$\frac{\partial L}{\partial \gamma_k} - \frac{\mathrm{d}}{\mathrm{d}t}\frac{\partial L}{\partial \dot\gamma_k} = \frac{\sqrt{\|\dot{\boldsymbol{r}}_1\|\|\dot{\boldsymbol{r}}_k\|}}{\dot\gamma_k}\left[\dot{\boldsymbol{r}}_1\cdot\boldsymbol{r}_k - \boldsymbol{r}_1\cdot\dot{\boldsymbol{r}}_k + \frac{1}{2}\boldsymbol{r}_1\cdot\boldsymbol{r}_k\left(\frac{\dot{\boldsymbol{r}}_1\cdot\ddot{\boldsymbol{r}}_1}{\|\dot{\boldsymbol{r}}_1\|^2} - \frac{\dot{\boldsymbol{r}}_k\cdot\ddot{\boldsymbol{r}}_k}{\|\dot{\boldsymbol{r}}_k\|^2}\right)\right].$$

Assuming $\frac{\sqrt{\|\dot{\boldsymbol{r}}_1\|\|\dot{\boldsymbol{r}}_k\|}}{\dot\gamma_k}$ is not zero at any point, we can divide with it, then the Euler-Lagrange equations to be solved are given with:

$$\dot{\boldsymbol{r}}_1\cdot\boldsymbol{r}_k - \boldsymbol{r}_1\cdot\dot{\boldsymbol{r}}_k + \frac{1}{2}\boldsymbol{r}_1\cdot\boldsymbol{r}_k\left(\mathit{\Gamma}_1 - \mathit{\Gamma}_k\right) = 0, \qquad k = 2,\ldots,n, \tag{19}$$

where 'Christoffel divergences' $\mathit{\Gamma}_i = \frac{\dot{\boldsymbol{r}}_i\cdot\ddot{\boldsymbol{r}}_i}{\|\dot{\boldsymbol{r}}_i\|^2}$, $i = 1,\ldots,n$ are introduced to simplify the equation.
